# Beige/brown fat-mediated cardiac protection from high-fat diet is dependent on adipocyte beta3-adrenergic receptor

**DOI:** 10.64898/2026.07.27.739389

**Authors:** L Cascarano, L Michel, H Esfahani, D De Mulder, A Melecchi, C Bouzin, A Loriot, J Ambroise, S Pezzica, F Carli, S Sabatini, L Gatto, C Dessy, A Gastaldelli, J-L Balligand

## Abstract

Cardiometabolic diseases associated with obesity are continuously increasing worldwide. Yet, current therapeutic strategies remain insufficient to improve patient outcomes. The beta3-adrenergic receptor (β3AR) promotes lipolysis in adipose tissue (AT) and thermogenesis specifically in brown adipose tissue (BAT). In mice, BAT activation also improves systemic metabolism and limits cardiometabolic damage. While BAT is limited in humans (e.g., with ageing and obesity), β3AR activation induces beige adipocytes within white adipose depots with similar thermogenic properties. To study the role of adipocyte β3AR in the regulation of cardiac metabolism and remodeling, mice with/without adipocyte-specific β3AR genetic deletion were fed a high-fat-sucrose (HF-S) diet and treated with the selective β3AR agonist CL316,243 (CL). The metabolic and cardiac protection following β3AR activation was abrogated upon β3AR deletion in adipocytes, together with the beiging of the epididymal (visceral) AT, highlighting a critical role of adipose β3AR signalling in mediating these benefits. Multi-omic analysis of AT and cardiac samples identified CL-induced secreted mediators of the crosstalk between AT and the heart. Therefore, adipocyte β3AR is critical for the adipose-cardiac communication and supports the therapeutic potential of targeting β3AR for the management of obesity-related cardiometabolic diseases.

## Introduction

Once regarded as a passive energy reservoir, adipose tissue (AT) is now recognized as a dynamic endocrine organ essential for metabolic homeostasis by regulating insulin sensitivity, immune responses and glucose metabolism (1,2). Through the secretion of adipokines, lipids, microRNAs (miRNAs) and other mediators, AT communicates with distant organs including the liver, skeletal muscle and the heart (3,4). This endocrine function is supported by the cellular heterogeneity of the tissue, which comprises not only adipocytes but also immune, stromal and vascular cells that together maintain tissue integrity (5).

Adipocytes can be classified into white, brown and beige cells, each with distinct metabolic roles. White adipocytes store lipids and regulate systemic energy availability, while brown adipocytes are highly oxidative and thermogenic via UCP1-dependent and independent uncoupling (6). In adult humans, small brown depots are present and remain responsive to cold or β3-adrenergic stimulation (7). Beige adipocytes, which are localised within white adipose tissue (WAT) and expand in response to sympathetic activation, combine features of both white and brown adipocytes (8).

The functional diversity of AT is profoundly influenced by its anatomical location (1). Subcutaneous adipose tissue (SAT), capable of healthy expansion and beiging, is generally protective against metabolic dysfunction (8,9). Conversely, upon excessive fat intake, visceral adipose tissue (VAT) evolves towards inflammation, hypoxia, increased fibrosis, extracellular matrix remodeling (ECM) and impaired adipogenesis (2,8). When the storage capacity of AT becomes saturated, excess lipids accumulate in ectopic organs such as the liver, skeletal muscle and heart, promoting insulin resistance, chronic inflammation and lipotoxicity (1,9,10).

Obesity is a chronic, multifactorial disease driven by genetic, environmental and behavioural factors, and is now recognized as a major contributor of cardiometabolic risk (9). Beyond the simple increase in body mass, obesity reflects a profound dysfunction of AT. In healthy conditions, AT can expand through adipogenesis (hyperplasia), allowing a preserved oxygen supply, limiting inflammation and maintaining insulin sensitivity (10). Conversely, impaired adipogenesis leads to excessive adipocyte hypertrophy, tissue hypoxia and ECM remodeling, which together promote fibrosis and chronic low-grade inflammation (11,12). These alterations compromise lipid storage capacity and disrupt adipokine secretion, with reduced adiponectin and increased leptin and inflammatory mediators (13).

These pathological changes support the well-established link between obesity and cardiovascular disease (CVD) (9,14,15). Obesity induces haemodynamic alterations, including plasma volume expansion and activation of the renin-angiotensin-aldosterone system (RAAS) with activation of the sympathetic nervous system (SNS) which drive structural remodeling of the heart (12). Left ventricular (LV) hypertrophy, impaired diastolic function and increased interstitial fibrosis and vascular stiffness are hallmarks of obesity-induced cardiovascular phenotype (16,17). Thus, dysfunctional AT, associated to altered endocrine profile, emerges as a central determinant of cardiometabolic risk (16). For instance, osteopontin, an adipokine secreted by the VAT of aged mice, has been shown to promote interstitial heart fibrosis (18).

In this context, targeting β3AR has attracted considerable interest. β3AR is G protein-coupled receptor (GPCR) predominantly expressed in adipocytes, where it plays a central role in regulating lipolysis and thermogenesis of AT (19). In white adipose depots, β3AR activation promotes beige adipocyte recruitment, while in BAT, it drives thermogenic gene expression and energy expenditure (20–22). The β3AR agonist mirabegron, at doses clinically used to treat overactive bladder disease (23), has been shown to stimulate BAT activity in humans, increase energy expenditure and improve lipid and glucose metabolism (24–29), although higher doses can induce cardiovascular side effects (25,27,28). Intriguingly, beige fat induction can improve the function of organs lacking β3AR expression, suggesting the existence of adipose-derived mediators with potential systemic, including cardiovascular effects (30).

Although epidemiological data suggest an association between active BAT and reduced prevalence of cardiometabolic and CVDs, the mechanisms linking thermogenic adipocytes (beige), β3AR activation and cardiovascular remodeling during obesity remain poorly understood (31). Supporting this hypothesis, a recent work has reported that AT beiging ameliorates diastolic dysfunction and improves lipid metabolism in a murine model of heart failure with preserved ejection fraction (HFpEF) (32).

In this study, we sought to determine whether the cardiometabolic benefits of β3AR activation depend specifically on adipocyte β3AR signalling. To address this question, we combined pharmacological stimulation with a genetic model of adipocyte-specific β3AR deletion and assessed the metabolic, adipose and cardiac consequences of chronic β3AR activation during diet-induced obesity. Using this approach, we were able to directly evaluate the contribution of adipocyte β3AR to metabolic regulation, AT remodeling and the progression of obesity-related cardiac remodeling.

## Material et Methods

### 1.1. Mice

Animals were housed in a 12:12-h day-night cycle with ad libitum access to water and chow diet (except when specified) in ventilated cages. All experiments were conducted in male mice to avoid potential confounding effects related to female hormonal fluctuations. This study was carried out in accordance with the NIH Guide for the Care and Use of Laboratory Animals and European Directive 2010/63/EU and was approved by local ethical committees. Body weight was recorded prior to diet initiation, monitored weekly during the first month, and then every two weeks until the end of the experimental period. Depending on the protocol, mice were fed for 4 to 6 months with a high-fat-sucrose (HF-S) diet (SAFE Diets, 59% fat and 27% carbohydrates, including 13% sucrose) before euthanasia and collection of adipose tissues and organs.

#### 1. C57Bl6/J mice +/- HF-S diet +/- CL316,243

36 male C57Bl6/J mice (9-week-old) were obtained from Janvier Labs (Le Genest Saint-Isle, France). Mice were randomized into four groups; 17 receiving a chow diet (SAFE Diets, Augy, France) and 19 were fed the HF-S diet. Within each group, half were implanted with a 4-week osmotic mini-pump delivering the β3AR agonist CL316,243 (Bio-techne, USA) immediately before diet initiation. A second CL mini-pump (diffusion period of 6 weeks) was implanted 4 weeks before sacrifice.

#### 2. AdipoCre^+/-^; *Adrb3*^-/-^ mice +/- HF-S diet

To generate transgenic mice with adipocyte-specific deletion of *Adrb3*, homozygous floxed *Adrb3* mice (in a C57Bl6/J background) were crossbred with heterozygous Adiponectin-Cre (AdipoCre^+/-^) or AdipoCre^-/-^ as littermate controls (wild-type) obtained from Jackson labs (AdipoCre-ERT, #025124). 9-week-old mice from both genotypes were randomly and equally divided into four groups. After 3 weeks of acclimatisation, two groups of AdipoCre^+/-^; *Adrb3*^-/-^were assigned to either a chow (CD) or HF-S diet as described above. To induce Cre recombinase activation, all mice received intraperitoneal injections of tamoxifen (30 mg/kg) for five consecutive days prior to diet initiation. One single dose injection was administered after 8 and 19 weeks to ensure efficient recombination in newly differentiated adipocytes.

#### 3. AdipoCre^+/-^; *Adrb3*^-/-^ mice + HF-S diet +/- CL316,243

In a separate cohort, 12-week-old AdipoCre^+/-^; *Adrb3*^-/-^ mice and wild-type AdipoCre^-/-^; *Adrb3*^-/-^were implanted with a 6-week osmotic mini-pump delivering the β3AR agonist (CL) immediately prior to the start of a 6-month HF-S diet. New 6-week mini-pumps were implanted at weeks 11 and 21 to ensure continuous agonist infusion throughout the entire feeding period.

### 1.2. Echocardiography

Mice were anesthetized by 1-3% isoflurane inhalation and were submitted to a two-dimensional echocardiography with a Vevo 2100 Imaging system (VisualSonics, Toronto, ON, Canada). The left ventricular (LV) parasternal long axis view was recorded in B-mode to measure LV internal volumes at end of diastole (LVEDv) and systole (LVESv), used to calculate the ejection fraction (EF; B-mode). M-mode view was performed to measure LV posterior wall (PW) and interventricular septum (IVS) thicknesses and LV internal dimensions (LVID), allowing the assessment of the fractional shortening (FS). LV mass was calculated as 1.053 X [(LVIDd + PWd + IVSd)^3^ – LVIDd^3^]. Measurements were carried out by the same operator. Echocardiographic tracings and analysis were performed in a blinded fashion using Vevo LAB software.

### 1.3. Glucose and insulin tolerance tests

Intraperitoneal glucose tolerance tests (IP-GTT) and insulin tolerance tests (IP-ITT) were performed at baseline and regularly during the diet period. For IP-GTT, mice were fasted for 6 h before intraperitoneal injection of glucose (2 g/kg body weight). Blood glucose levels were measured from tail blood at 0, 15, 30, 60, and 120 min using a Contour Next glucometer (Ascensia). For IP-ITT, mice were fasted for 4 h prior to injection of insulin (0.5 U/kg body weight), and glucose levels were measured using the same protocol. In compliance with European regulations and local ethics guidelines, blood sampling was performed by microsampling from a 1-mm tail tip incision. For each time point, a drop of blood from the gently milked tail was applied directly to the test strip without restraining the animal. Serial samples for ITT and GTT (performed 24 h apart) were obtained by removing the scab from the previous sampling site. Results are expressed as plasma glucose concentration (mg/dL) for GTT and as percentage of basal glycemia for ITT.

### 1.4. Tissue Collection and Dissection

At the time of euthanasia, adipose depots were carefully dissected and collected, including inguinal (subcutaneous), epididymal (visceral), brown, and pericardial adipose tissues. For each mouse, the right inguinal and epididymal pads were consistently sampled.

To arrest cardiac contraction in diastole, hearts were immersed in 50 mM KCl solution immediately after excision. The heart was then weighed, and the right ventricle removed to isolate the left ventricle (LV) for further analysis. Portions of LV, inguinal, and epididymal adipose tissues were fixed in formalin and embedded in paraffin for histological studies.

### 1.5. RNA Extraction and Real-Time Reverse Transcription-Polymerase Chain Reaction

Total RNA was isolated from epididymal, inguinal and brown adipose tissues using TRI Reagent (Fermentas) and homogenized with Precellys Evolution (Bertin Technologies). Extraction was performed according to standard protocols. Residual genomic DNA was eliminated by DNase treatment (RQ1 RNase-free DNase, Promega). RNA from heart and pericardial fat was extracted using Maxwell and specific kit (Maxwell® RSC simplyRNA Tissue Kit, A1340, Promega). Subsequently, 1 microgram of total RNA was reverse transcribed into cDNA using RevertAid Reverse Transcriptase with oligo(dT) and random hexamer primers (Thermo Fisher Scientific). Quantitative PCR was performed with low ROX Takyon (Eurogentec) on 1:25 diluted cDNA with the CFX96 real-time system (Bio-Rad Laboratories, Vienna, Austria). Relative gene expression was calculated using the 2^–ΔΔCt (Livak method) method and normalized to housekeeping gene expression [hypoxanthine phosphoribosyl transferase (HPRT)]. The exon-overlapping amplicons were amplified using the following calibrated primer sets:

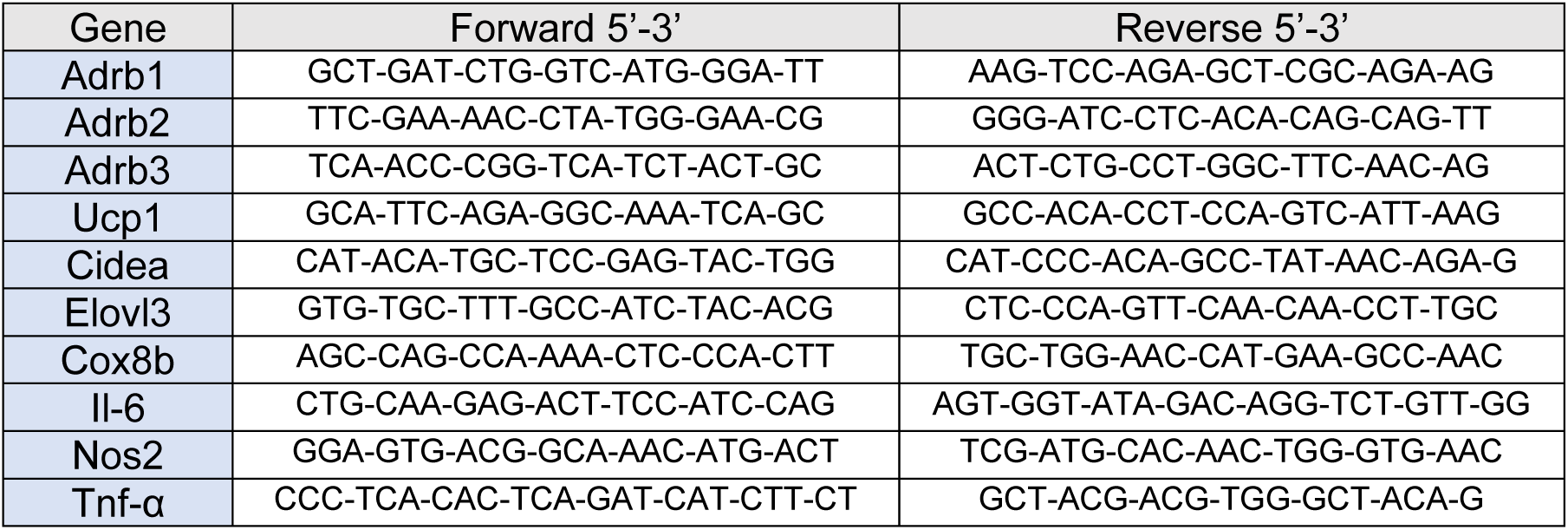

### 1.6. Transcriptomic analysis

RNA of heart and epididymal adipose tissue from C57Bl/6J mice lean or obese, treated with CL or not (N=3 per group) was extracted using Maxwell and specific kit (Maxwell® RSC simplyRNA Tissue Kit, A1340, Promega). After RNA integrity was evaluated using an Agilent DNR-472 (15 nt) HS RNA kit on a 5200 Fragment Analyzer to ensure RIN value greater than 7, 100ng were processed according to the standard protocol for the Watchmaker RNA library prep kit. Sequencing was performed on Illumina NovaSeq 6000, generating 60M reads (30M PE) 2×150 bp. Fastq files were processed using a standard RNAseq pipeline including Trimmomatic (v0.39) (32) to remove low quality reads, HISAT2 (v2.2.1) (33) to align reads to the mouse genome (GRCm38), and gene expression levels were evaluated using feature Counts from Subread (v2.0.3) (34) and Mus_musculus.GRCm38.94.gtf. Differential expression analyses were performed with DESeq2 Bioconductor package v1.50.2 (35). Genes encoding secreted proteins were identified using annotations from the Universal Protein Resource (UniProt) database. Gene set enrichment analyses were done using clusterProfiler v4.18.3 (36) on gene sets from the MSigDB database.

### 1.7. Metabolomics

After thoracotomy, hearts were rapidly freeze-clamped, weighed and stored in liquid nitrogen. Polar metabolites were extracted as previously described (33). Briefly, after tissue homogenization (Precellys Evolution Homogenizer, Bertin Instruments, Frankfurt, Germany) in cold methanol, proteins were precipitated by centrifugation (25 minutes at 13300 rpm and 4°C). Subsequently, polar metabolites were isolated using a modified Folch extraction (34), with chloroform: methanol (2:1, v/v) and water (33,34). Amino acids, organic acids and glycolytic intermediates were quantified from the polar (upper) phase, by adding labelled ^13^C ^15^N internal standards (20ul,100uM MSK-CAA-1 Canonical Amino Acid Mix standard, CIL Cambridge, MA, USA; 20ul, 25uM MSK-OA-1 Labeled Organic Acid Mix, CIL Cam bridge, MA, USA), together with xylose (1ul, 666uM, Merck KGaA, Darmstadt, Germany) and norleucine (5ul, 1525uM, Sigma-Aldrich, St. Louis, MO, USA) before homogenization. Polar phases were then dried under gentle nitrogen flux and were derivatized using 10 µl of Methoxyamine hydrochloride (Merck KGaA, Darmstadt, Germany) solution in pyridine (20 mg/mL) for 30 minutes at 60°C, followed by 50ul of N, O-bis(trimethylsilyl) trifluoroacetamide with 1% trimethylchlorosilane (TMCS) (Merck KGaA, Darmstadt, Germany) and 50ul acetonitrile for 1hour at 60°C and then analysed by gas chromatography mass spectrometry. The untargeted acquisitions were performed with a single quadrupole (GC 7890A/MS 5975C, Agilent, Santa Clara, CA) equipped with a capillary column (DB-5MS J&W, l 30 m; i.d. 0.25 mm; film thickness 0.25 μ m, J&W, Agilent) and the identification of the metabolites was performed using the Fiehn library. The targeted acquisition was performed with a gas chromatography coupled with tandem mass spectrometry triple quadrupole (GC 8890/ MS-TQ 7000D, Agilent, Santa Clara, CA) equipped with a capillary column (HP-5Q J&W, l 30 m; i.d. 0.25 mm; film thickness 0.25 J&W, Agilent) and the quantification was done using the labelled internal standards as previously described (33).

### 1.8. Western Blot

Pieces of brown adipose tissues were lysed in RIPA buffer containing protease and phosphatase inhibitor cocktail (P8340, 4 906 837 001 Merck) using Precellys Evolution (Bertin technologies) before being centrifuged 15 minutes at 20000 rcf to remove fat supernatant. Proteins were quantified using Pierce BCA protein assay (Thermo Fisher Scientific). Samples were prepared in Laemmli solution and heated for 10 min at 70°C. 40 µg of proteins were loaded and separated by SDS-PAGE and transferred onto PVDF membranes. Membranes were blocked 1 hour with 5% milk in TTBS (Tris buffered saline with 0,1% Tween 20, Sigma), and were incubated with specific primary antibodies (B3AR anti-rabbit Abcam, #94506, 1/1000 and HSP90 anti-rabbit C45G5, 1/5000) overnight at 4°C. After three washing in TTBS, membranes were incubated by a secondary antibody for 1 hour (Anti-rabbit, 1/5000) in 1% milk. Membranes were washed three times in TTBS and signals were revealed with a chemiluminescent reagent (ECL, Amersham, Cytiva) on X-ray films in a dark chamber.

### 1.9. Fibrosis assessment

Left ventricular (LV) as well as inguinal and epididymal adipose tissue samples were fixed for 24h in 4% formaldehyde, embedded in paraffin and sectioned at 5 µm. Haematoxylin-eosin (HE) and picrosirius red (PSR) staining were performed using standard procedures.

For the assessment of fibrosis, PSR-stained LV and AT were digitized using a 3DHISTECH ScanII slide scanner, and computer-assisted analysis was performed with the image analysis tool Author version 2017.2 (Visiopharm, Hørsholm, Denmark). Total tissue section was delineated. Artifacts, pericardium and fascia surrounding AT were manually excluded before analysis. The interstitial fibrotic area was expressed as a percentage of stained area per total tissue area. For each heart, the mean fibrosis value was calculated from three independent LV sections.

### 1.10. Immunohistofluorescence

Paraffin sections were placed at 37°C for 24h prior to immunostaining. After deparaffinization, sections were processed according to the protocol previously described by Aboubakar et al. (35). Endogenous peroxidase activity was quenched with 3% hydrogen peroxide in isopropanol for 20 min. Heat-induced epitope retrieval (HIER) was then performed, followed by blocking in Tris-buffered saline (TBS) containing and 0.1% Tween-20 (TBS-T) and 5% bovine serum albumin (BSA). The first primary antibody was incubated 1 h at room temperature in TBS-T supplemented with 1% BSA, followed by 40 min incubation with horseradish peroxidase (HRP)-conjugated polymer secondary antibodies (Agilent, Cat. No. K4003). Detection was performed using tyramide signal amplification (TSA) with CF-conjugated tyramides (diluted 1000x). Sequential labeling of up to three additional markers was achieved by repeating HIER and TSA cycles as indicated in Table 2. For LV sections, wheat germ agglutinin (WGA)-rhodamine (RL-1022, Vector Laboratories, dilution 1:50) was used to delineate cardiac myocyte membranes. After final washes in TBS-T (or PBS for WGA), nuclei were counterstained with Hoechst 33342 (Sigma, Cat. No. 1453) diluted in distilled water and slides were mounted using Dako fluorescence mounting medium (Agilent, Cat. No. S3023). Slides were scanned using a Zeiss Axioscan.z1 at X20 magnification.

### 1.11. Macrophage infiltration analysis

F4/80, CD68 and CD206 immunostainings were quantified on entire tissue sections using Author version 2017.2. On each slide, tissue sections were automatically surrounded at low magnification. Delineations were visually checked and manually corrected if required. Cells were detected at high resolution (X20) with a nuclear-based cell segmentation relying on the Hoechst staining. Cells were then classified according to the F4/80, CD86 and CD206 staining intensities above fixed thresholds. The same parameters were kept constant for all slides. Results are expressed as percentages of stained cells.

### 1.12. Cell area measurement

Perilipin-1, haematoxylin-eosin and WGA staining were used to automatically delineate each adipocyte or cardiac myocyte, respectively, and to measure cell area (expressed in µm^2^).

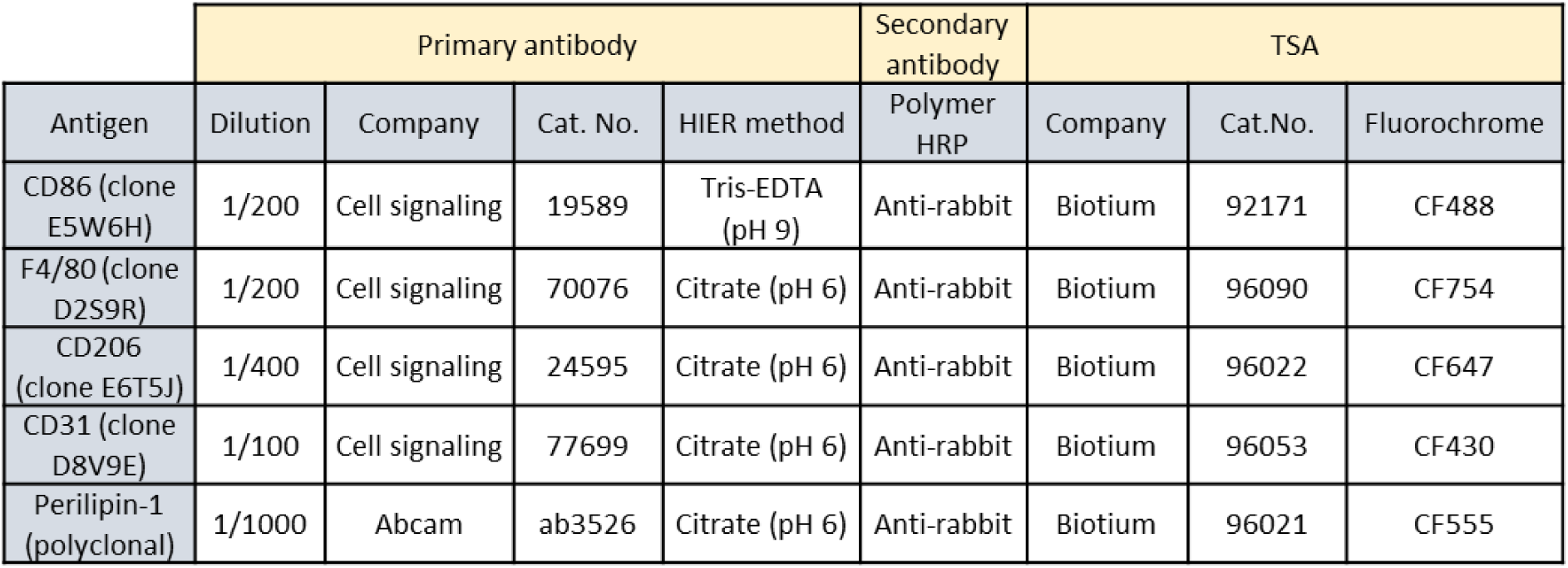

### 1.13. Statistical Analysis

Data are presented as means ± SEM. Statistical comparisons were performed using unpaired t-tests, one-way or two-way ANOVA, or two-way ANOVA for repeated measures, as appropriate. When normality assumptions were not met, nonparametric tests (Kruskal-Wallis or Mann-Whitney) were applied. A p-value < 0.05 was considered statistically significant. Post hoc analyses following ANOVA were performed using Sidák’s multiple-comparison correction. Statistical analyses and data visualization were conducted with GraphPad Prism (GraphPad Software, San Diego, CA, USA).

### 1.14. NicheNet prediction and prioritization of putative messengers

Although initially developed for single-cell transcriptomic data analysis, the nichenetr R package (v2.2.1.1) was applied to predict which ligands among the differentially expressed genes (DEGs) in adipose tissue are most likely to regulate the DEGs observed in heart tissue. This prioritization was achieved using a pre-computed ligand-target matrix, which integrates prior public biological knowledge to score regulatory potentials. Finally, the circlize R package (v0.4.17) was utilized to visualize the predicted regulatory links between these key adipose-derived ligands and their cardiac target genes in a circular chord diagram.

### 1.15. Genome-Scale Metabolic Modelling and integration of gene expression data

Genome-scale metabolic modelling (GSMM) was employed to integrate transcriptomics data within a metabolic network framework and to relate gene-expression alterations to metabolomic changes. Genome-scale metabolic models (GEMs) provide curated stoichiometric representations of cellular metabolism, linking genes, enzymes, reactions, metabolites, and metabolic pathways that can be used for systematic interpretation of omics data at the reaction and pathway levels (36). In this study, differential cardiac gene-expression ratios between experimental groups were mapped onto gene-protein-reaction (GPR) rules in the generic mouse GEM Mouse1 (37) to infer reaction ratio scores. While gene expression is not expected to quantitatively predict reaction flux across all genes, prior studies have shown that condition-specific changes in gene expression can provide a proxy for relative shifts in metabolic flux distributions (38,39). To calculate the reaction ratio scores we followed the methodology proposed by Fang *et al.* (39) and by Tsouka et al (40): for reactions associated with a single gene, the corresponding expression ratio was assigned directly to the reaction; reactions controlled by multiple genes, protein complexes, represented by “AND” relationships, were resolved using the geometric mean of the relevant gene-expression ratios, whereas “OR” relationships, representing isoenzymes, were resolved using the arithmetic mean. Composite GPR rules were decomposed sequentially according to the same logic. The resulting reaction ratio scores were then used to interrogate the GEM and to identify pathways associated with transcriptional alterations induced by HFD and potentially reversed by CL treatment. Where metabolomics data were available, detected metabolites were mapped to their associated reactions and subsystems and integrated with the corresponding reaction-level ratio scores, providing complementary evidence to support the interpretation of pathway-level metabolic remodeling.

## Results

### Systemic β3AR activation limits weight gain, improves metabolic profile and protects the heart against adverse remodeling in obese mice

To investigate the systemic effects of β3AR activation in obesity, male C57Bl6/J mice were fed either a chow diet (CD) or a high-fat-sucrose (HF-S) diet for 26 weeks and were treated or not with the β3AR agonist CL316,243 (CL) (Figure 1A). This mouse model under obesogenic diet previously characterized by us (41) displays a reproducible cardiac structural remodeling in response to HF-S feeding. Importantly, we verified that *Adrb3* is prominently expressed in adipose tissue (AT) of these mice, but that the receptor is undetectable in cardiac tissue (suppl. Figure 1A), in line with previous demonstration in mouse cardiac myocytes (42). CL treatment was delivered via an osmotic mini-pump implanted at the beginning and during the last month of diet. Although CL is highly specific for the mouse β3AR, the dosage was adjusted in preliminary experiments to avoid off-target effects on cardiac β1AR, as confirmed from unaltered mean blood pressure by implanted telemetry in awake mice (suppl. Figure 1B). As expected, mice fed the HF-S diet showed a marked increase in body weight compared with CD controls. CL treatment transiently attenuated weight gain during the infusion periods, with a significant reduction in body weight observed at the end of each CL treatment phase (Figure 1B).

**Figure 1:**
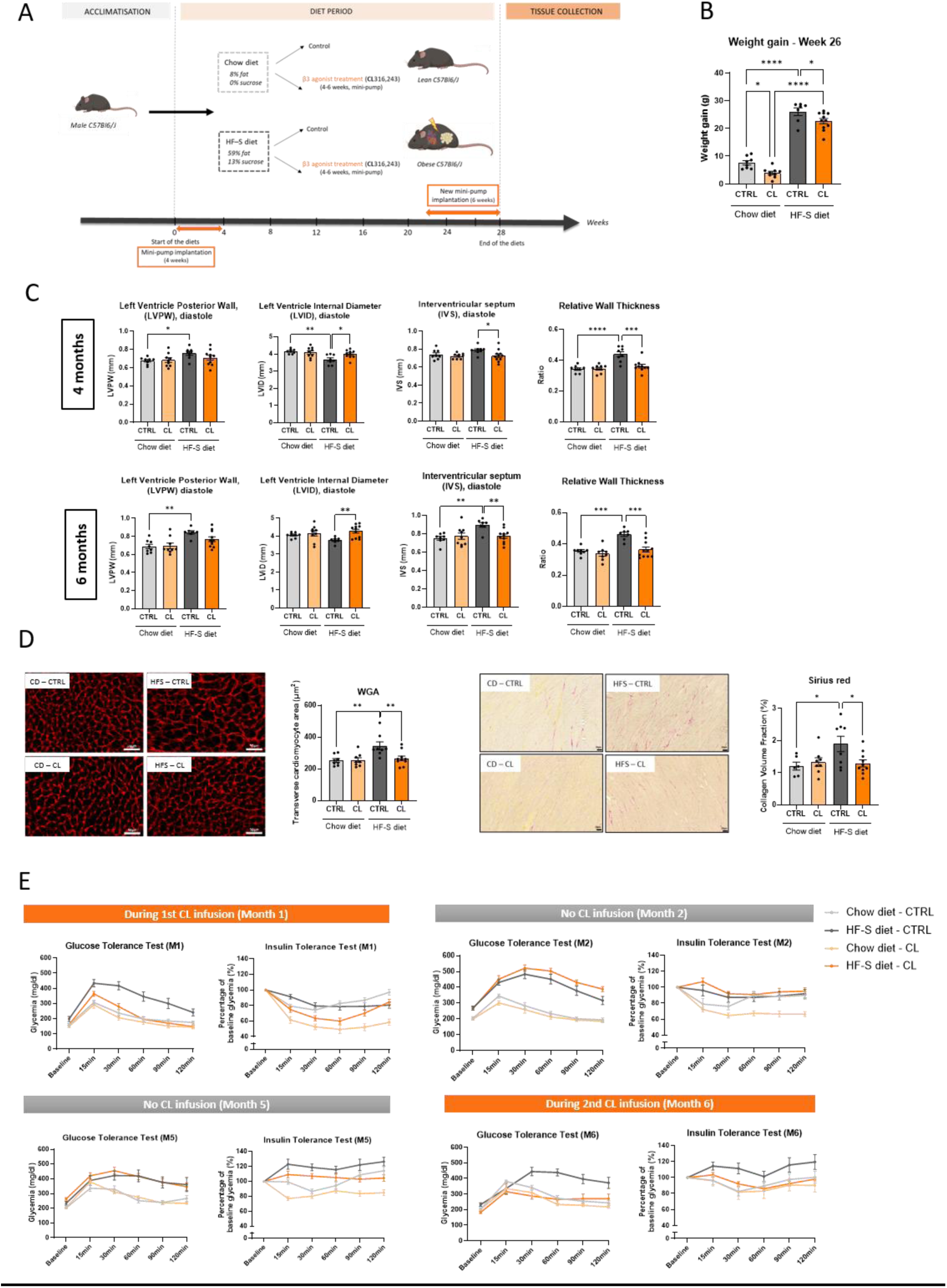
Metabolic and cardiac characterization of C57BL/6J mice subjected to chow or high-fat-sucrose diets with or without β3AR agonist CL316,243. (A) Schematic representation of experimental design; (B) Body weight gain after 26 weeks of diet; (C) Echocardiographic assessment of left ventricle posterior wall (LVPW), interventricular septum (IVS), left ventricle internal diameter (LVID) in diastole and relative wall thickness (RWT) measured after 4 or 6 months of diet; (D) Quantification of left ventricular cardiac myocyte cross-sectional area by wheat germ agglutinin (WGA) staining and of myocardial interstitial fibrosis by Picrosirius red staining at the end of diet (scale bars = 50 µm); (E) Intraperitoneal glucose tolerance test (GTT, 2g/kg) and insulin tolerance test (ITT, 0.5U/kg) during (months 1 and 6), after (month 2) and before (month 5) CL316,243 (CL) treatment (orange) on mice submitted to a chow (light) or a HF-S (dark) diet. Data are presented as means +/- SEM (N = 7-11 per group) and analysed by 2-way ANOVA (*p<0,05, **p<0,01, ***p< 0,001, ****p<0,0001).

Echocardiographic analysis performed after 4 and 6 months of diet exposure revealed significant cardiac remodeling in HF-S mice. Specifically, these mice displayed increased left ventricular posterior wall (LVPW) thickness, interventricular septum (IVS) thickness, and relative wall thickness (RWT), along with a decrease in left ventricular internal diameter (LVID), reflecting concentric hypertrophy. Interestingly, CL-treated HF-S mice did not exhibit these structural alterations, suggesting a protective effect of β3AR activation on cardiac remodeling. Notably, this protection persisted even after discontinuation of CL treatment (month 4) (Figure 1C).

Histological analysis confirmed the echocardiographic data. Wheat germ agglutinin (WGA) staining revealed cardiac myocyte hypertrophy in HF-S mice, which was prevented by CL treatment. Picrosirius red (PSR) staining showed increased interstitial fibrosis in the heart of obese mice, whereas CL-treated animals exhibited significantly reduced fibrotic areas (Figure 1D).

To assess glucose homeostasis, glucose tolerance (GTT) and insulin tolerance (ITT) tests were performed at different time points throughout the diet period. HF-S-fed mice developed impaired glucose tolerance and insulin resistance as early as the first month of diet. CL treatment prevented these metabolic alterations during the infusion period, maintaining glucose and insulin sensitivity at levels comparable to CD mice (month 1). However, this effect was lost once CL infusion stopped (months 2 and 5), indicating a transient metabolic improvement. The second CL infusion at the end of the diet restored glucose tolerance and insulin sensitivity (AUC: 1916 ± 117.5 *vs* 1365 ± 124.7 in CL-treated obese mice, *p*<0.001), showing a β3AR agonist-mediated correction effect on metabolic parameters (Figure 1E).

Together, these results demonstrate that systemic β3AR activation durably prevents obesity-induced cardiac hypertrophy and fibrosis, despite transient metabolic improvement.

### β3AR activation induces beiging selectively in epididymal adipose tissue of obese mice

We next examined the effects of CL treatment on AT biology. As expected, HF-S-fed mice exhibited a marked increase in AT mass across all depots, including ING AT, EPI AT, BAT and pericardial fat. In contrast, CL administration selectively reduced the mass of brown and pericardial depots in obese mice (Figure 2A). Similarly, CL treatment decreases the weight of retroperitoneal and subcutaneous interscapular AT, whereas no major changes was observed in perirenal and perivascular AT (suppl. Figure 2 and suppl. Table 1). These morphological changes were accompanied by alterations in the expression of thermogenic and beige markers (Ucp1, Cidea, Cox8b, and Elovl3), with variabilities across adipose depots. Notably, CL treatment increased the expression of these genes in obese mice specifically in the EPI AT depot, without significant changes detected in the ING AT or pericardial AT (Figure 2B). Moreover, CL induced a shift toward smaller adipocytes in EPI AT and to a lesser extent in ING AT (suppl. Figure 3).

**Figure 2:**
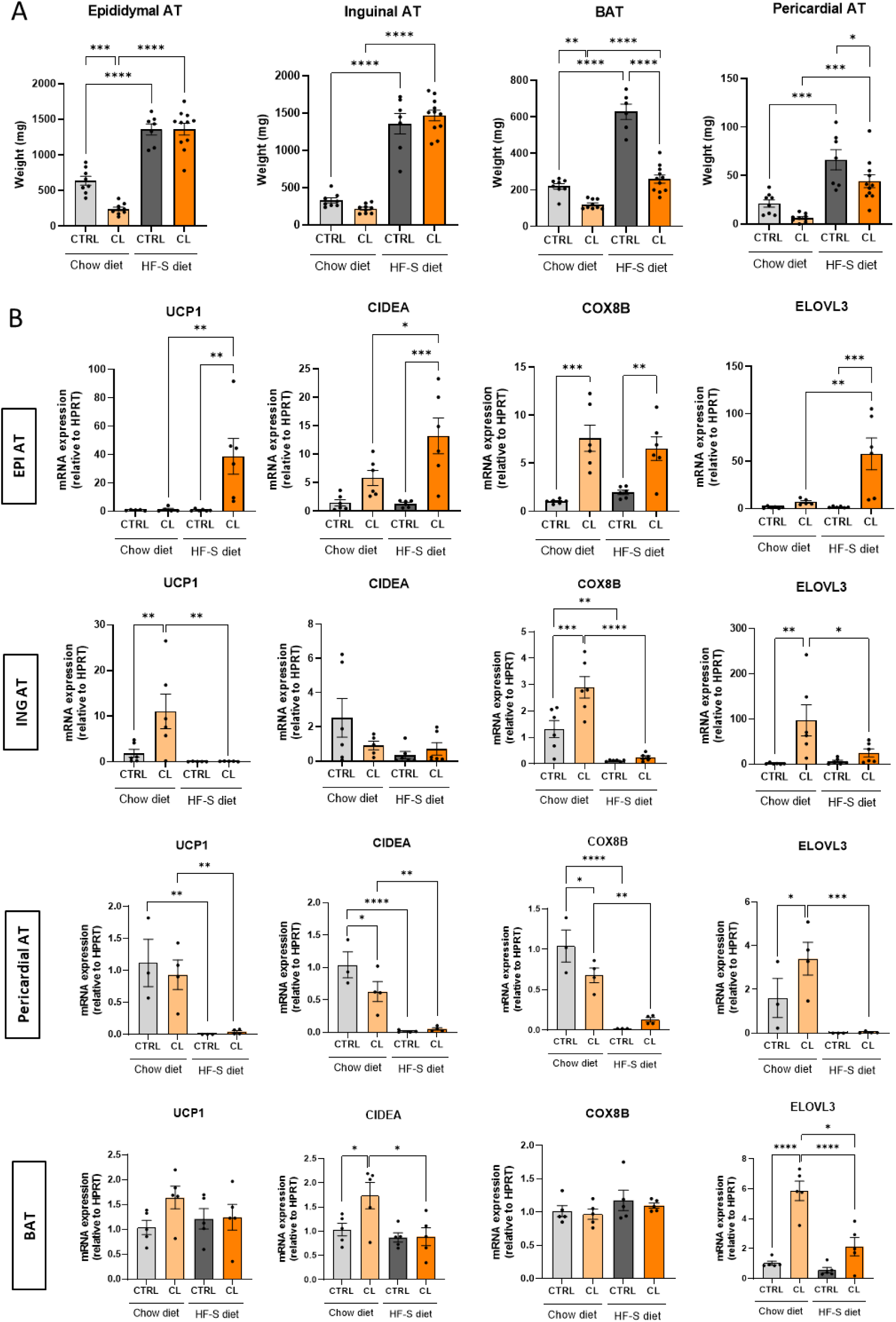
Adipose tissue characterisation in chow- and HF-S-fed mice treated or not with CL316,243. (A) Weight of epididymal, inguinal, brown and pericardial adipose tissue (AT) depots after 26 weeks of diet; (B) Relative mRNA expression of thermogenic genes *Ucp1*, *Cidea*, *Cox8b*, and *Elovl3* in epididymal (EPI), inguinal (ING), pericardial AT and brown AT (BAT). Data are presented as means +/- SEM (N = 7-11 per group) and analysed by 2-way ANOVA (*p<0,05, **p<0,01, ***p< 0,001, ****p<0,0001).

We carried out both molecular and histological characterisation of inflammation parameters in EPI, ING and brown AT under chow and HF-S diets. No clear pattern appeared under obesogenic diet except an increase in Il-6 and Nos2 in BAT of obese mice that was corrected by CL treatment (suppl. Figure 4A). Moreover, in EPI AT, the obesogenic diet seemed to increase a specific subset of macrophages (CD86+/CD206+), that was normalized with CL. This was not apparent in ING AT (suppl. Figure 4B).

Collectively, these findings indicate that systemic β3AR activation by CL316,243 differentially modulates AT remodeling across depots, with selective activation of beiging markers in EPI AT of obese mice.

### Conditional adipocyte-specific β3AR deletion *per se* does not induce systemic metabolic alterations or modify heart structure

To determine whether the cardioprotective effects of β3AR activation were mediated through its expression in adipocytes, we generated an adipocyte-specific β3AR knockout model using the tamoxifen-inducible Cre-lox system. Mice carrying floxed *Adrb3* alleles were crossed with Adiponectin-CreERT2 mice to obtain AdipoCre^⁺/-^/*Adrb3*^fl/fl^ (referred to as B3 adipoKO) and littermate control AdipoCre^⁻/⁻^/*Adrb3*^fl/fl^ (WT) mice (Figure 3A).

**Figure 3:**
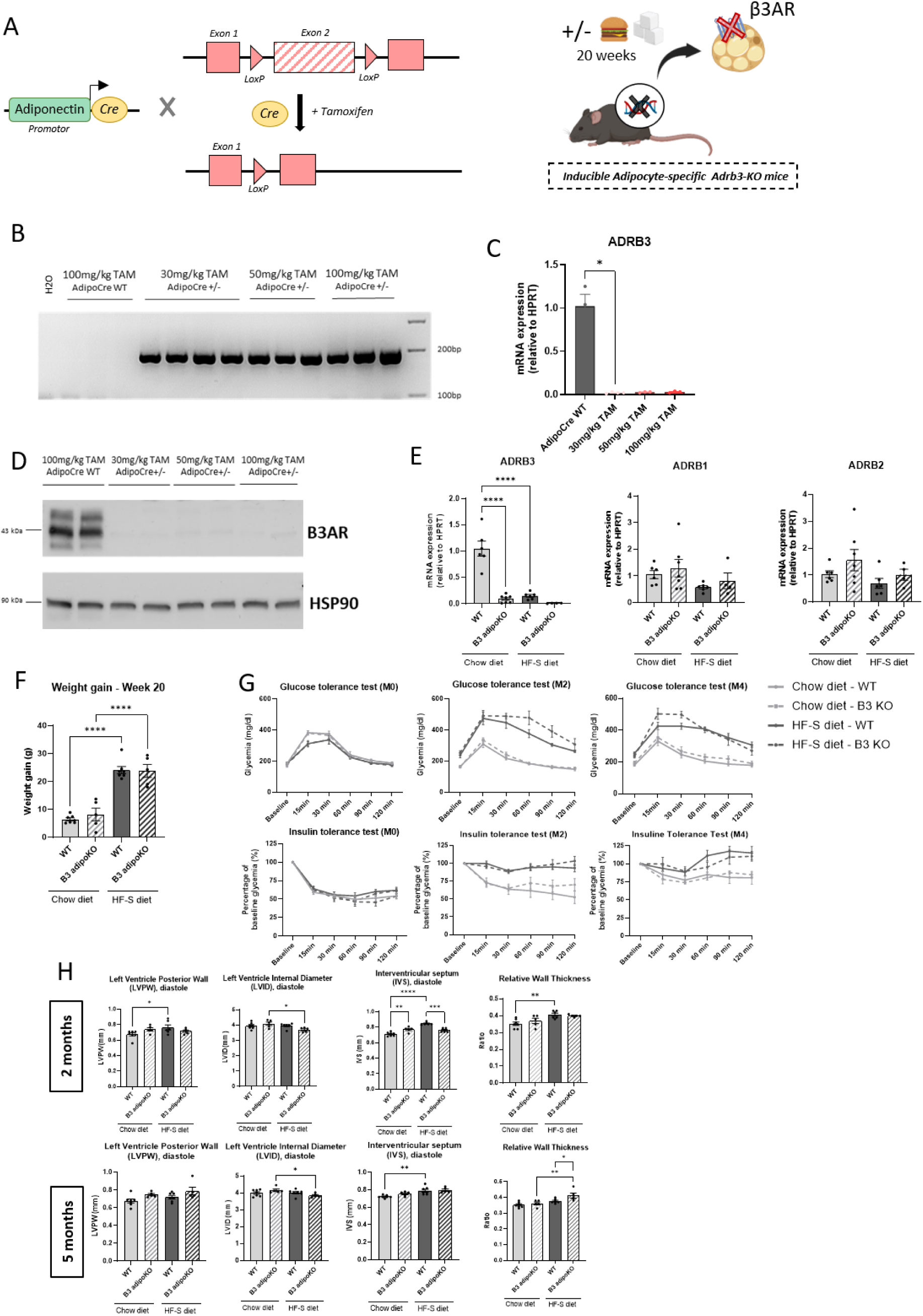
Generation and characterization of inducible adipocyte-specific β3AR knockout mice (AdipoCre^+/-^;*Adrb*3^fl/fl^). (A) Schematic representation of the inducible *Adrb3* knockout model. Tamoxifen (TAM) administration activates Cre recombinase under the adiponectin promoter, excising floxed *Adrb3* alleles and inducing adipocyte-specific β3AR deletion. Transgenic and WT mice were fed chow or HF-S diets for 5 months. Validation of recombination efficiency two days after intraperitoneal tamoxifen injection (for 5 consecutive days): (B) PCR detection on epididymal AT genomic DNA; (C) *Adrb3* mRNA expression in isolated adipocytes of inguinal AT and; (D) β3AR protein levels in brown AT by western blot. Negative controls (AdipoCre−/−, WT) were treated with 100 mg/kg TAM and AdipoCre positive (AdipoCre+/−) mice received 30, 50, or 100 mg/kg TAM (N = 3-4 per group); (E) Relative mRNA expression of *Adrb3*, *Adrb2*, and *Adrb1* in isolated inguinal adipocytes; (F) Body weight gain after 20 weeks of diet; (G) Intraperitoneal glucose tolerance test (GTT, 2g/kg) and insulin tolerance test (ITT, 0.5U/kg) of at baseline (M0), 2 months (M2) and 6 months (M6) of diet of WT (solid line) and B3 adipoKO (hatched) mice submitted to a chow (light) or a HF-S (dark) diets; (H) Echocardiographic evaluation of left ventricle posterior wall (LVPW), interventricular septum (IVS), left ventricle internal diameter (LVID) in diastole and relative wall thickness (RWT) at 2 and 5 months of diet. Data are presented as means +/- SEM (N = 6-9 per group) and analysed by 2-way ANOVA (*p<0,05, **p<0,01, ***p< 0,001, ****p<0,0001).

Tamoxifen (TAM) administration at doses of 30, 50 or 100 mg/kg efficiently induced recombination of the floxed *Adrb3* gene, as confirmed by PCR on genomic DNA on EPI AT (Figure 3B) and by a remarkable reduction in *Adrb3* mRNA expression (Figure 3C) in isolated adipocytes from ING AT. Consistent with the mRNA data, β3AR protein levels were undetectable in BAT by western blot after TAM treatment (Figure 3D). In contrast, *Adrb1* and *Adrb2* mRNA expression remained unchanged, confirming the specificity of the deletion (Figure 3E).

Following tamoxifen induction, mice were maintained on either chow or HF-S diets for 20 weeks. No differences in body weight gain, glucose or insulin tolerance were observed between B3 adipoKO and WT mice, under CD or HF-S diet (Figure 3F-G), indicating that β3AR loss in adipocytes did not markedly affect systemic metabolism.

Echocardiographic analysis did not identify consistent changes between WT and β3AR adipoKO mice under either diet regimen at 2 or 5 months, except an increase in relative LV wall thickness at 5 months in β3AR adipoKO (Figure 3H), suggesting that adipocyte-specific deletion of β3AR alone may not be sufficient to promote significant cardiac remodeling under basal conditions (i.e., in absence of CL stimulation).

Consistent with body weight evolution, adipocyte β3AR deletion did not significantly affect total AT mass, except for a reduction in pericardial fat observed in obese mice (Figure 4A). Morphometric analysis of adipocyte size distribution revealed a tendency for larger adipocytes in B3 adipo KO mice under both CD and HF-S diets, more marked in epididymal than inguinal depots (Figure 4B).

**Figure 4:**
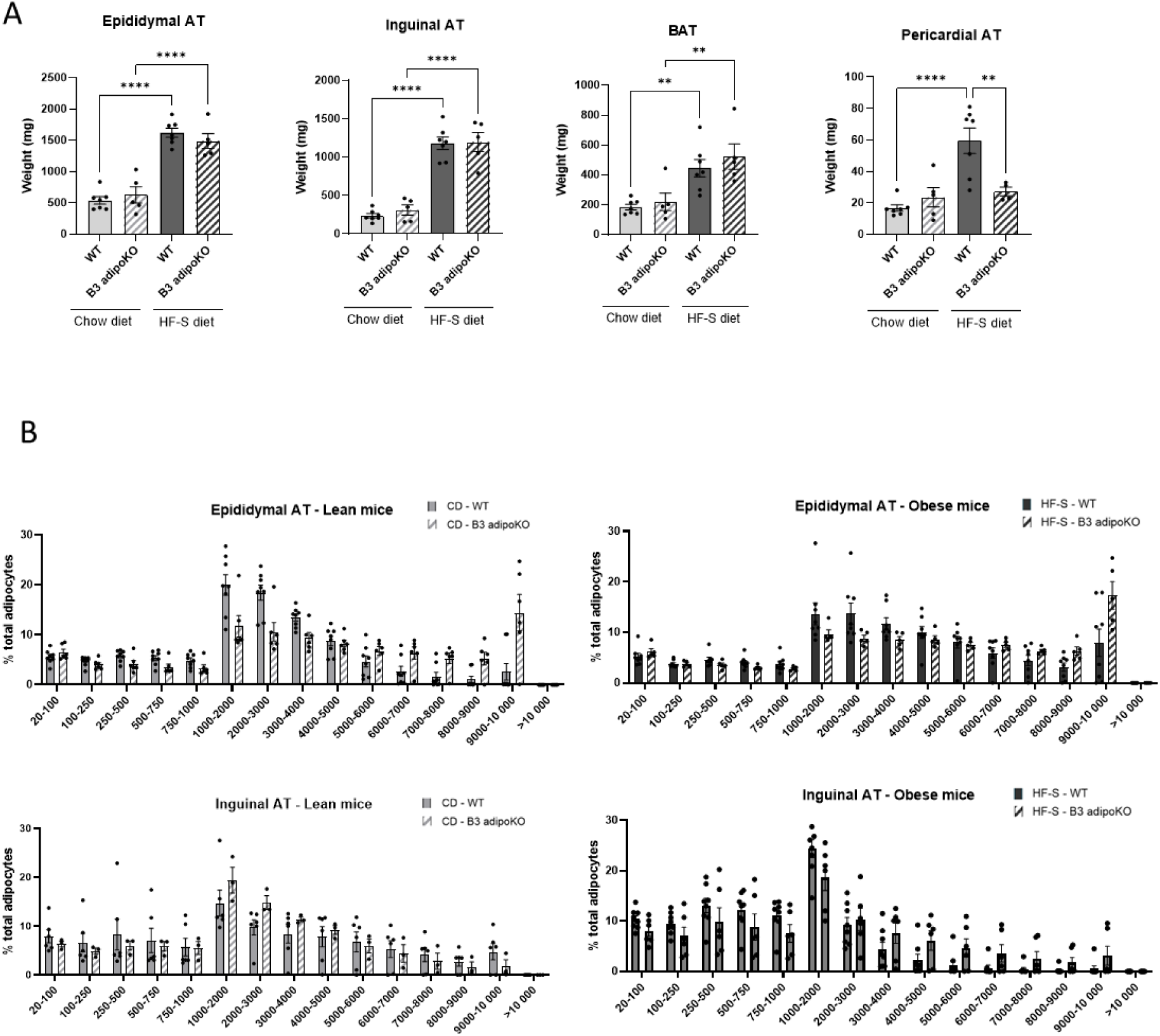
Adipose tissue characterisation in chow- and HF-S-fed *Adrb3* transgenic and WT mice. (A) Weight of epididymal, inguinal, brown and pericardial AT after 20 weeks of diet; (B) Adipocyte size distribution (20-1000 µm^2^) in epididymal (upper) and inguinal (lower) AT in B3 adipoKO (hatched) or wild-type (WT) mice submitted to a chow (CD, light grey) or a high-fat-sucrose (HF-S, dark grey) diet. Data are presented as means +/- SEM (N = 5-9 per group) and analysed by 2-way ANOVA (**p<0,01, ****p<0,0001).

### Loss of adipocyte β3AR abrogates the cardioprotective effects of CL

To directly assess whether the cardioprotective effects of β3AR activation depend on its expression in adipocytes, WT and B3 adipoKO mice were submitted to the HF-S diet for 26 weeks and were treated or not with CL throughout the feeding period (Figure 5A).

**Figure 5:**
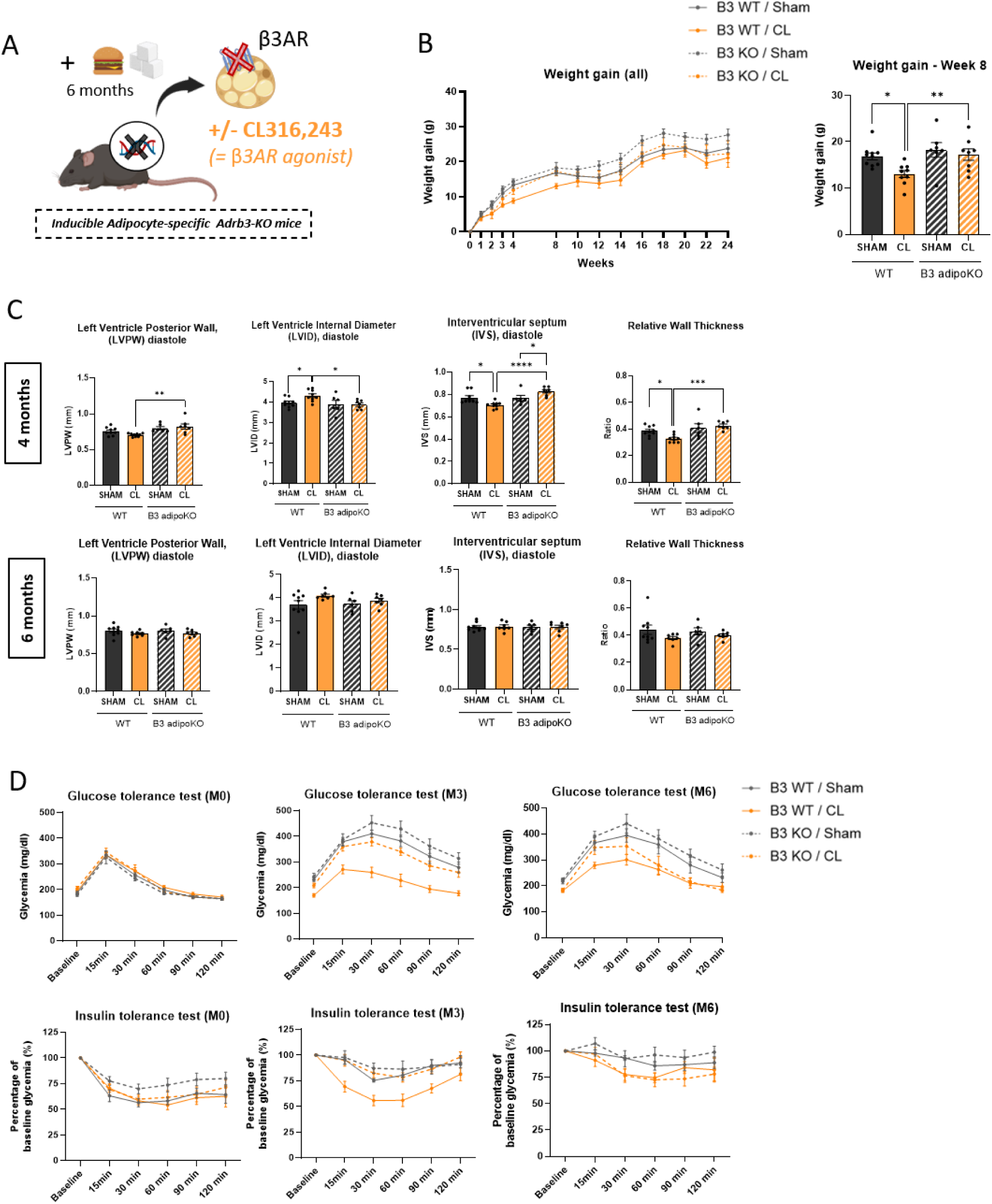
Metabolic and cardiac characterization of adipocyte-specific β3AR knockout (B3 adipoKO) and WT mice fed a high-fat-sucrose diet with or without β3AR agonist CL316,243 treatment. (A) Schematic illustration of the experimental design; (B) Body weight evolution (from baseline to week 24) of WT (solid line) and B3 adipoKO (dotted line) mice, treated with (orange) or without (grey) CL316,243 (left) and weight gain at week 8 (right); (C) Echocardiographic assessment of left ventricle posterior wall (LVPW), interventricular septum (IVS), left ventricle internal diameter (LVID) in diastole and relative wall thickness (RWT) measured after 4 or 6 months of diet; (D) Intraperitoneal glucose tolerance test (GTT, 2g/kg) and insulin tolerance test (ITT, 0.5U/kg) at baseline (M0), 3 months (M2) and 6 months (M6) of diet. Data are presented as means +/- SEM (N = 7-10 per group) and analysed by 2-way ANOVA (*p<0,05, **p<0,01, ***p< 0,001, ****p<0,0001).

As previously observed, CL treatment initially reduced body weight gain in HF-S-fed WT mice compared with untreated controls, as illustrated by a significant difference at 8 weeks (13± 0,8g *vs* 16,9± 0,7g, *p*<0,05). This effect was completely lost in HF-S-fed B3 adipoKO mice (17,2±1,3g, *p*<0,01), which displayed similar weight gain under CL compared with their untreated controls. Notably, the CL-induced reduction in body weight gain in HF-S-fed WT mice was progressively attenuated over time (Figure 5B).

Echocardiographic assessment at 4 months indicated that HF-S-fed WT mice treated with CL exhibited reduced wall thickness and maintained normal LV morphometry, in contrast to their untreated HF-S controls, confirming the protective effect of β3AR activation on cardiac remodeling. Notably, this CL protection was completely abolished in B3 adipoKO mice, which developed concentric hypertrophy comparable to untreated HF-S mice. At a later time point, the cardiac protection by CL was lost even in WT mice (month 6) (Figure 5C).

At the metabolic level, CL treatment preserved glucose tolerance and insulin sensitivity in WT mice after three months of HF-S feeding, whereas these protective effects were absent in B3 adipoKO mice. However, as for cardiac effects, CL protection seemed to disappear with time and chronic exposure (Figure 5D).

Overall, these data confirmed the major role of the adipocyte β3AR in the β3AR agonist-mediated cardioprotection in obese mice.

### Adipocyte β3AR deletion prevents CL-induced beiging of epididymal AT

We next examined whether the lack of cardioprotection in B3 adipoKO mice was associated with altered AT phenotype. At the end of the HF-S diet, β3AR deletion had little impact on overall AT mass, with or without CL. As observed before, CL induced a significant reduction in pericardial fat mass in WT mice, but not in B3 adipoKO mice (Figure 6A).

**Figure 6:**
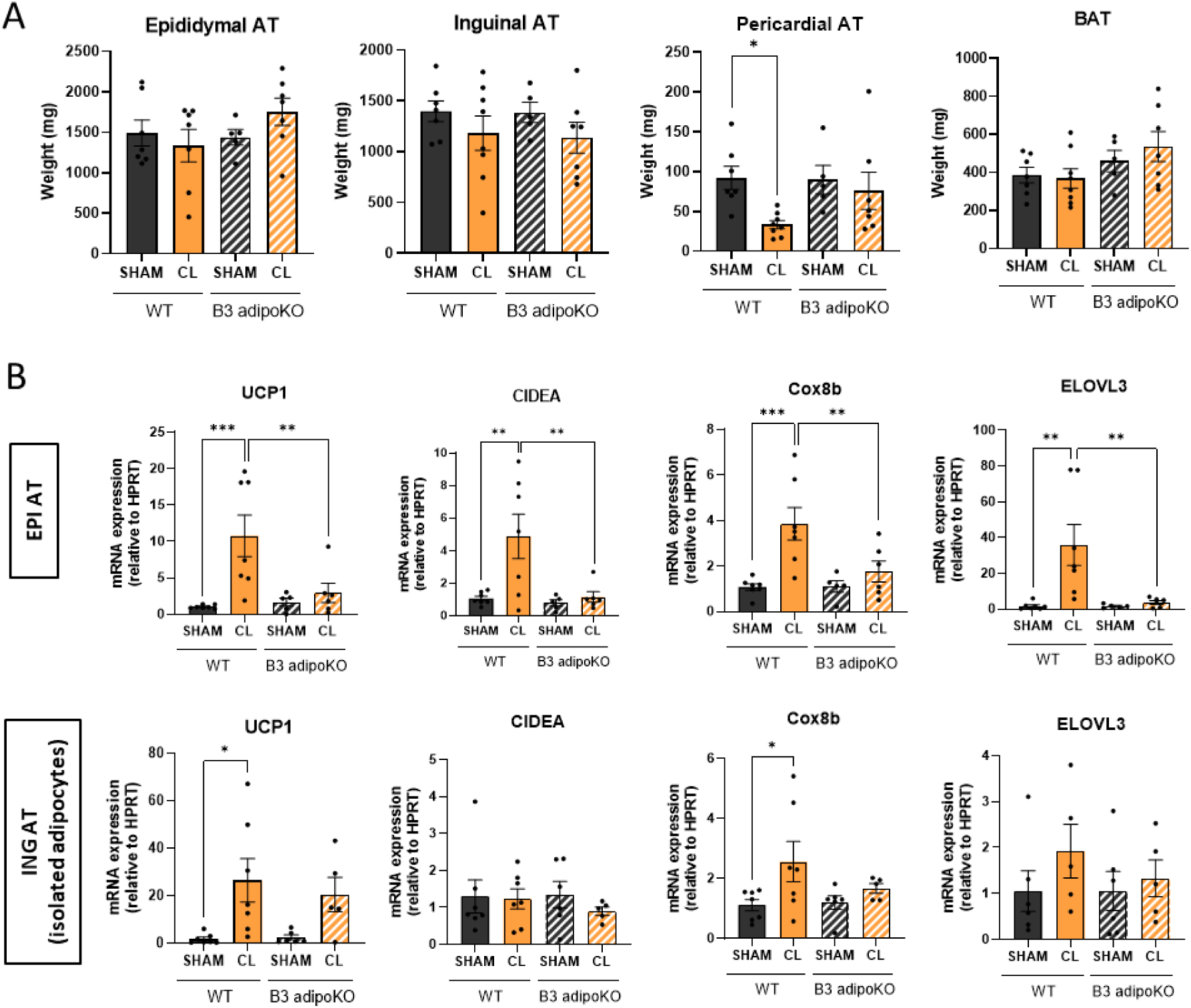
Adipose tissue characterisation in adipocyte-specific β3AR knockout (B3 adipoKO) or WT mice subjected to high-fat-sucrose diet with or without β3AR agonist CL316,243 treatment. (A) Weight of epididymal, inguinal, brown and pericardial AT after 24 weeks of diet; (B) Relative mRNA expression of thermogenic genes Ucp1, Cidea, Cox8b, and Elovl3 in epididymal (EPI) and isolated adipocytes from inguinal (ING) AT. Data are presented as means +/- SEM (N = 6-9 per group) and analysed by 2-way ANOVA (*p<0,05, **p<0,01, ***p< 0,001, ****p<0,0001).

In WT mice, CL treatment induced the expression of thermogenic markers (Ucp1, Cidea, Cox8b, Elovl3) specifically in EPI AT and in subcutaneous depots. This beiging response was completely abolished in B3 adipoKO mice, only in EPI AT, revealing that adipocyte β3AR expression is essential for CL-induced thermogenic activation in EPI AT of obese mice (Figure 6B).

Altogether, these results demonstrate that the early cardioprotective and metabolic benefits of β3AR activation depend on the presence of adipocyte β3AR. The absence of adipocyte β3AR abolishes both the induction of beige adipocytes in epididymal depots and the associated cardiometabolic protection, although the latter disappears upon chronic CL treatment.

### Integrated cardiac and adipose transcriptomic analysis identify potential pathways and candidate mediators involved in adipose-heart crosstalk

Having established a role of adipocyte β3AR in mediating CL-induced cardioprotection, we next sought to uncover the molecular mechanisms underlying this AT-heart communication. To this end, we performed unbiased transcriptomic analysis on both cardiac and epididymal AT using RNA sequencing.

Firstly, we analysed EPI AT from obese mice to characterize transcriptional changes induced by β3AR activation. Gene set enrichment analysis revealed that CL treatment significantly modulated several biological pathways compared to untreated HF-S mice. Interestingly, pathways related to inflammation (IFNϓ, IL-6/Jak/STAT3, TNF-α signalling), angiogenesis, hypoxia and epithelial-mesenchymal transition (EMT) were significantly downregulated. In contrast, oxidative phosphorylation, adipogenesis and FA metabolism were upregulated (Figure 7A-B).

**Figure 7:**
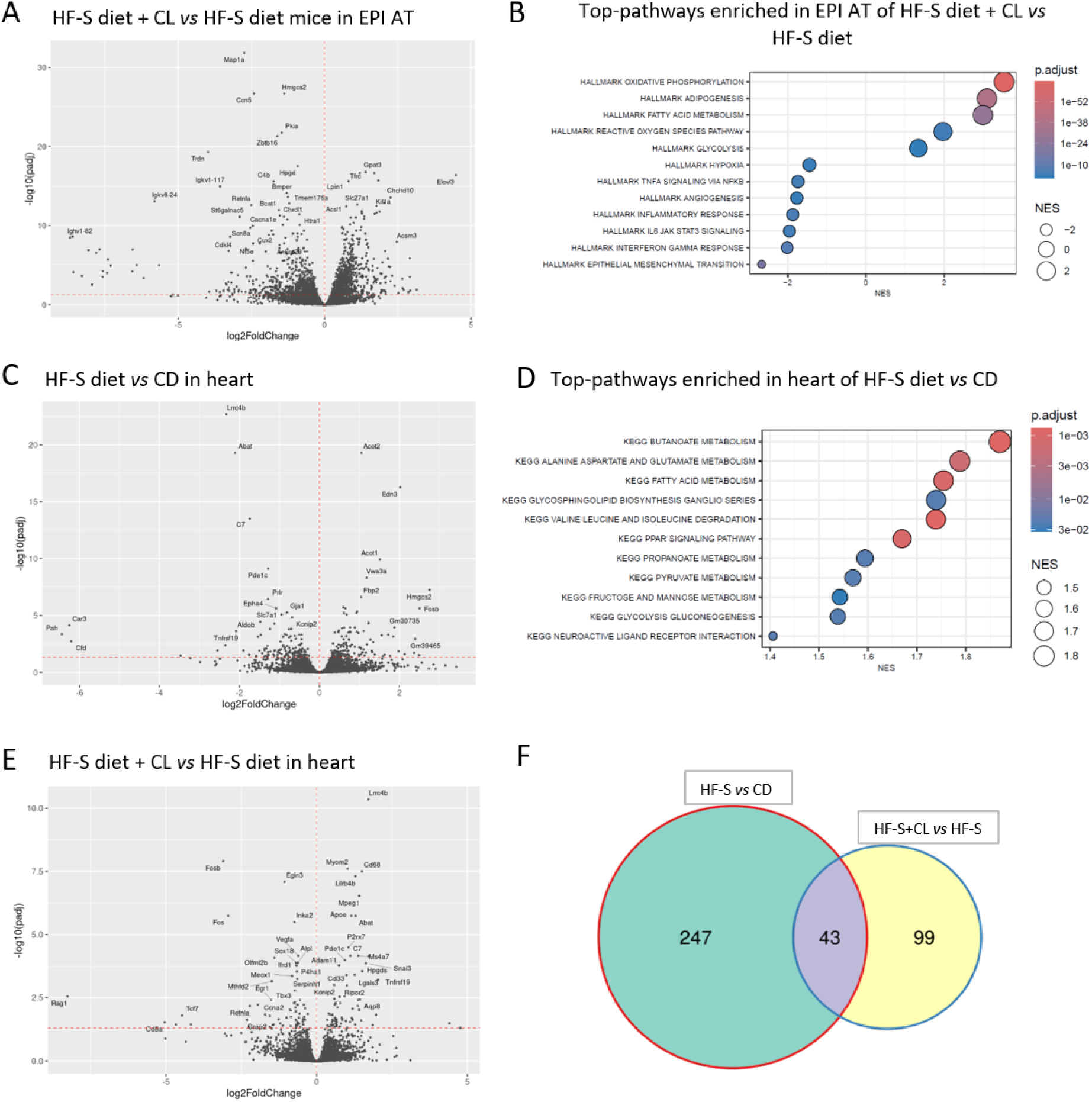
RNA sequencing analysis in epididymal adipose tissue and heart in chow- and high-fat-sucrose-fed mice treated or not with CL316,243. (A) Volcano plot of differentially expressed genes (DEGs) and; (B) Dot plot of 12 enriched hallmark pathways identified by gene set enrichment analysis (GSEA) in epididymal adipose tissue (EPI AT) of CL-treated high-fat-sucrose (HF-S) diet mice compared with untreated HF-S diet mice; (C) Volcano plot of DEGs and; (D) Dot plot of 11 enriched KEGG pathways in hearts from HF-S diet compared with chow diet (CD) mice; (E) Volcano plot of DEGs in hearts from HF-S mice treated with CL compared to untreated HF-S mice; (F) Venn diagram illustrating the overlap of 43 genes between DEGs identified in hearts from HF-S *vs* CD diet mice and CL-treated HF-S mice *vs* untreated HF-S mice, highlighting commonly regulated genes (N = 3 per group).

We next examined whether β3-adrenergic activation influences cardiac gene expression in the context of obesity by performing RNA sequencing analysis on cardiac tissue from the same mice.

We first compared hearts from CD and HF-S diet mice in the absence of CL treatment to determine the impact of diet-induced obesity on the cardiac transcriptome. This analysis identified 290 differentially expressed genes, reflecting the impact of obesity on the cardiac gene expression (Figure 7C). Functional enrichment analysis revealed that these genes were predominantly associated with metabolic processes including lipid metabolism (fatty acid metabolism and PPAR signalling, butanoate and propanoate metabolism) and glucose metabolism (glycolysis, gluconeogenesis, pyruvate, fructose and mannose) as well as amino-acid-related pathways (alanine, aspartate and glutamate metabolism; valine, leucine and isoleucine degradation) (Figure 7D).

We then evaluated the effect of CL treatment in HF-S-fed mice and identified 142 genes differentially expressed in heart between untreated and CL-treated HF-S mice, which encompass pathways related to immune and inflammatory responses, metabolic processes and extracellular matrix remodeling (Figure 7E). To determine whether CL reversed obesity-induced transcriptional alterations, we intersected the two datasets using a Venn diagram analysis (Figure 7F). This comparison revealed 43 genes that were both significantly altered by the HF-S diet and modulated in the opposite direction by CL treatment, indicating a transcriptional “rescue” signature (suppl. Table 2). These 43 genes highlighted pathways related to hypertrophic, fibrotic and hypoxic signalling (*Fos, Fosb, Egr1, Pdgfd, Egln3*) and cardiac signalling and electrophysiological regulation (*Kcnip2, Pde1c and Stk39*). In addition, several genes related to mitochondrial and metabolic processes (*Abat, Pcx, Slc25a13, Pah*) were also present in the gene set. Notably, the upregulation of stress and hypertrophy-associated genes induced by obesity was normalized by CL treatment, whereas several genes involved in mitochondrial and metabolic functions that were suppressed by HF-S feeding were restored upon AT β3AR activation.

### GSMM identifies metabolic alterations reversed by CL treatment

Based on DEG in hearts under HF-S +/-CL, we used Genome-Scale Metabolic Models (GSMM) to infer potentially altered metabolic pathways. By including all known biochemical reactions catalysed by enzymes and transporters encoded in the genome, GSMMs analyse and predict the behaviour of complex metabolic networks. Going from pathways to putative enzymes and individual transcripts, a normalized score is associated to the corresponding metabolic reaction(s), with scores above 1 indicating a likelihood of enhanced metabolic flux through this reaction.

This approach identified the upregulation of a number of genes and associated enzymatic reactions in CL-treated HF-S hearts, including those highlighted in Figure 8, that were associated with pyruvate metabolism (Pcx), glutamate and amino acid metabolism (e.g., Kyat, Abat, Pah, and glutamate transporter, Slc25a13), as well as 1-carbon metabolism (Mthfd2). These reactions were then integrated in metabolic pathways, as illustrated in Figures 9 and 10.

**Figure 8:**
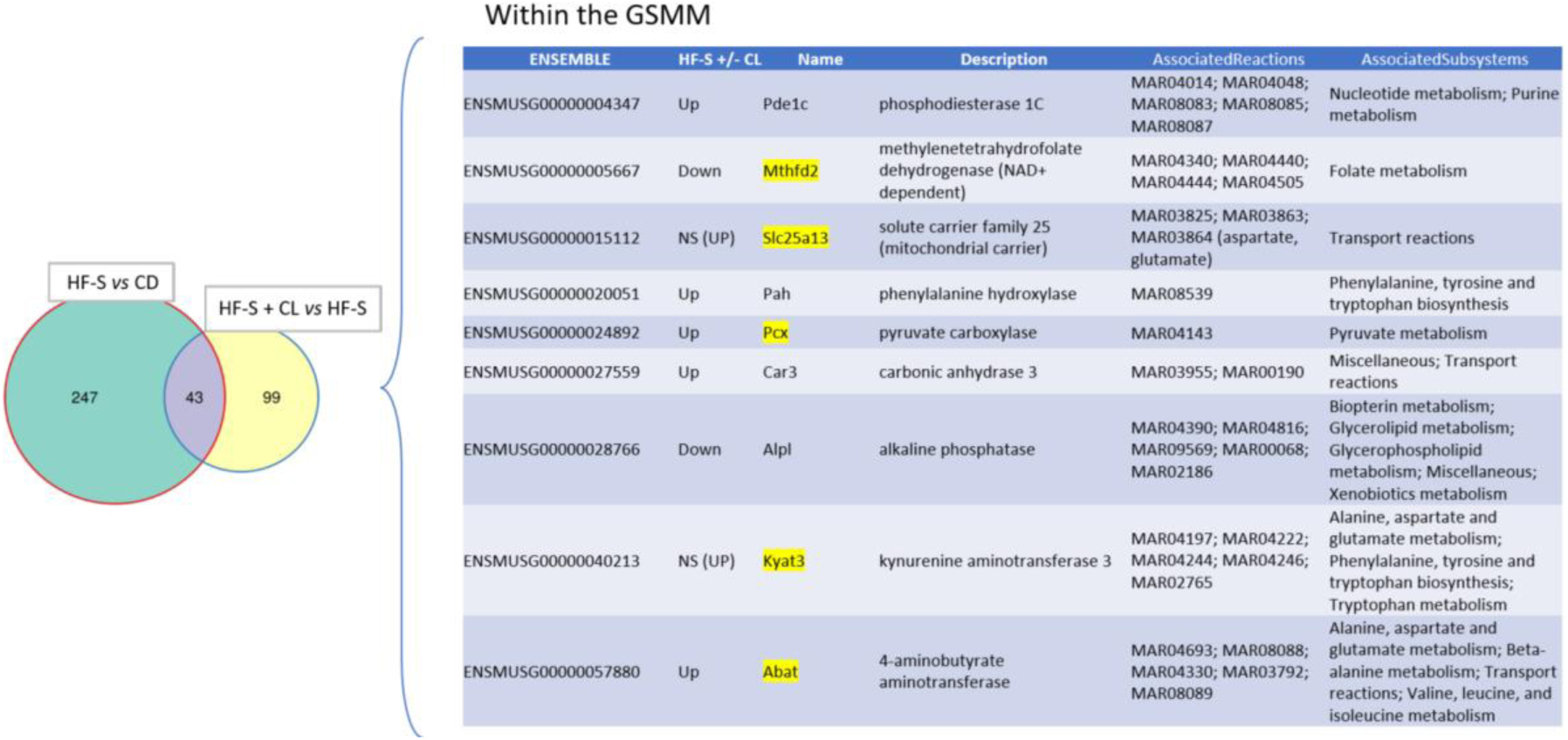
Metabolic pathways, corresponding enzymatic reactions and enzyme-coding genes identified in the mouse generic genome-scale model from the 43 intersected DEG between HF-S and HF-S+CL hearts. Highlighted are the gene names corresponding to enzymes involved in reactions modified under CL treatment, as illustrated in the model in Figure 9.

**Figure 9:**
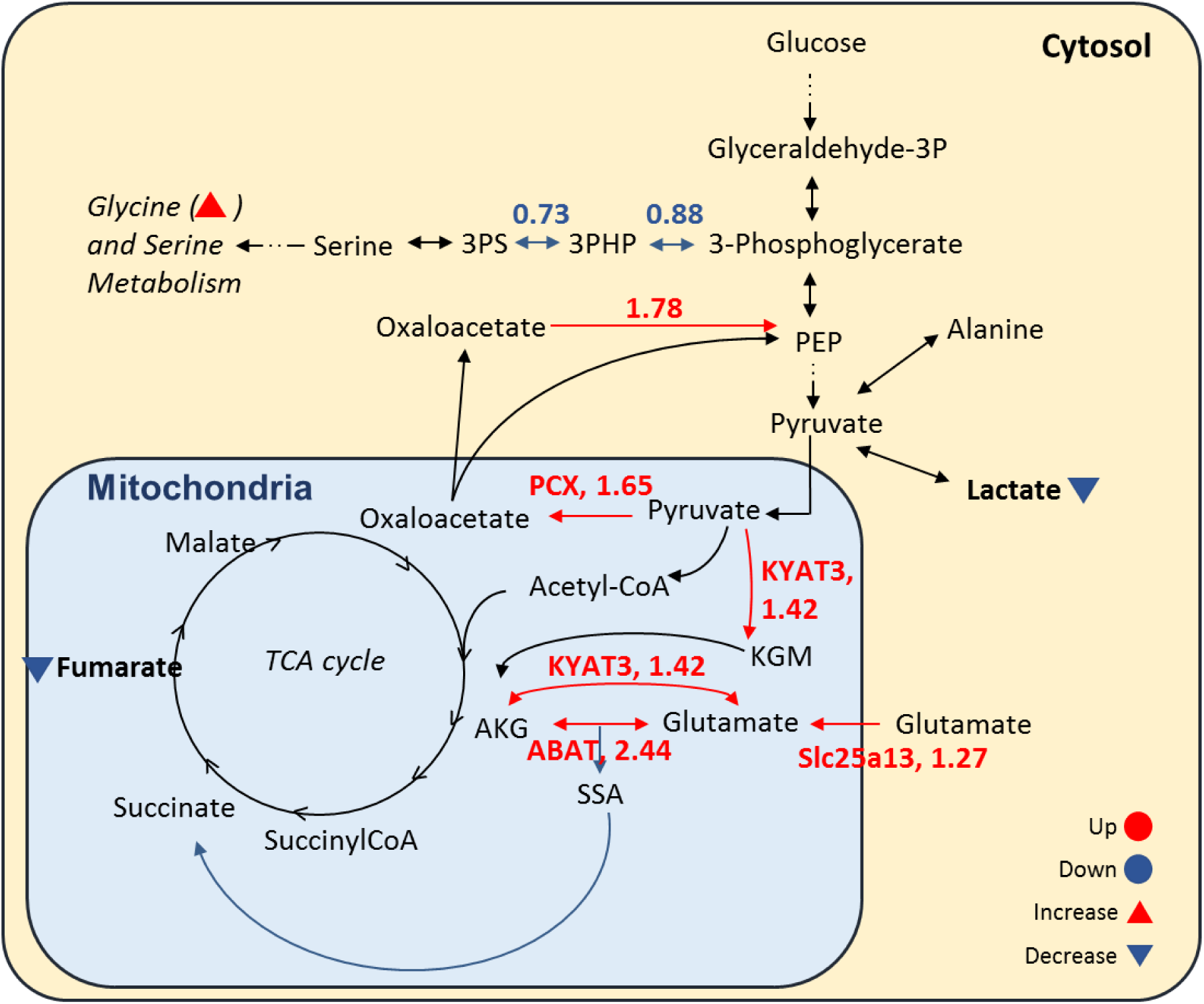
Metabolic network illustrating pathways alterations observed in the glycolysis, tricarboxylic acid (TCA) cycle, and pyruvate metabolism subsystems in HF-S conditions with *vs.* without CL. Reactions exhibiting changes in flux ratios greater than 1.25, lower than 0.9, or associated with differentially expressed genes are highlighted in red (upregulated) or blue (downregulated). Triangles indicate significant alterations in metabolite levels in HF-S+CL relative to HF-S alone, as observed in our metabolomic measurements, with red and blue color and upward and downward orientations denoting increases and decreases, respectively.

**Figure 10:**
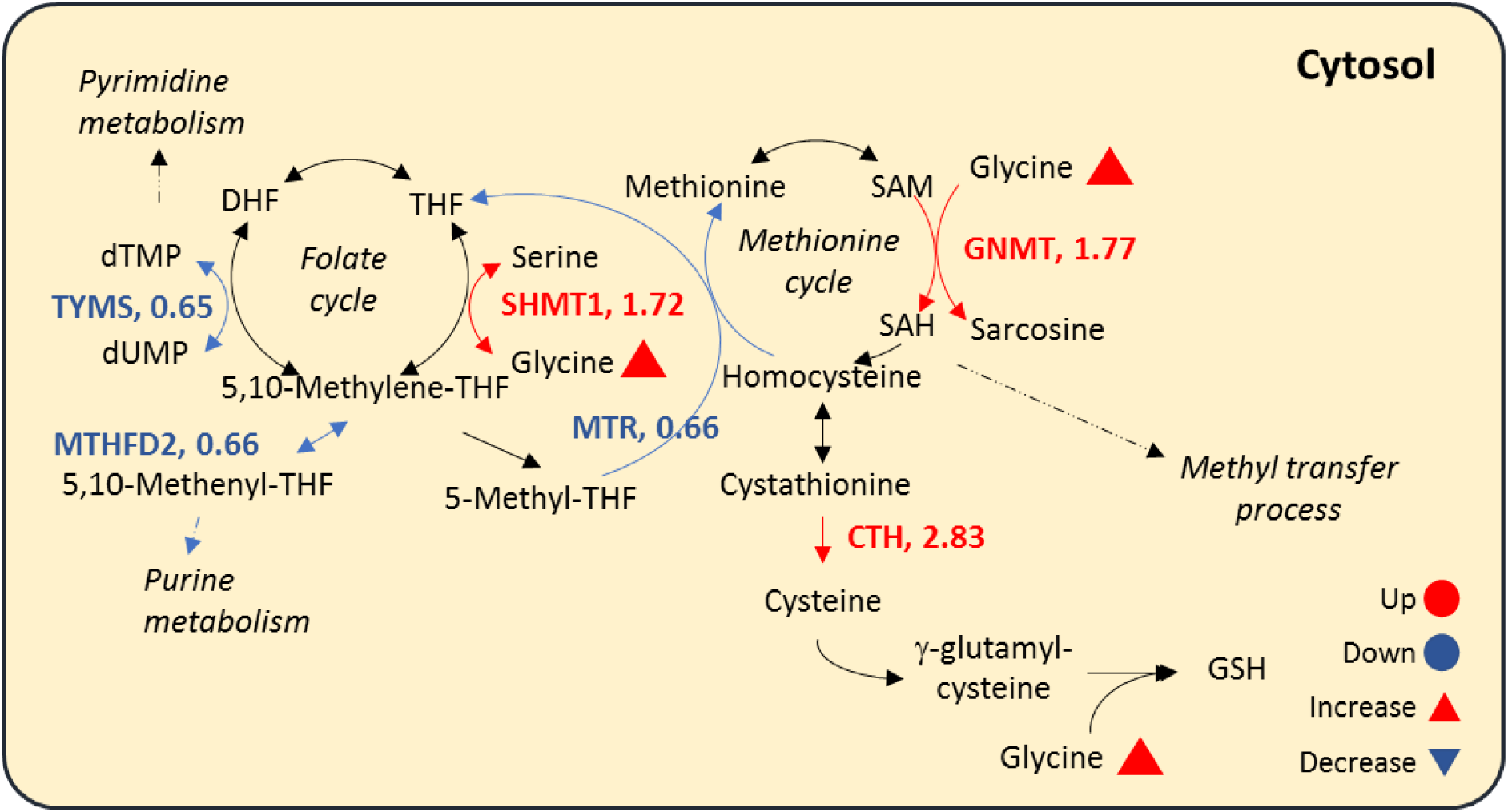
Metabolic network illustrating pathway alterations observed in the one carbon metabolism in HF-S conditions with versus without CL. Reactions exhibiting changes in flux ratios greater than 1.25, lower than 0.9, or associated with differentially expressed genes are highlighted in red (upregulated) or blue (downregulated). Triangles indicate significant alterations in metabolite levels in HF-S+CL relative to HF-S alone, as observed in our metabolomic measurements, with red and blue color and upward and downward orientations denoting increases and decreases, respectively.

Collectively, the positive scores (>1) indicate increases in key reactions metabolizing pyruvate and glutamate that would replenish the Krebs cycle. Notably, the model also predicted significant changes in the metabolism of lactate that resulted in a decrease in lactate accumulation as also shown by metabolomic analyses of heart tissues (see Figure 11 and suppl Figure 5). In addition, it would predict a downregulation of Serine production from glycolytic intermediates.

**Figure 11:**
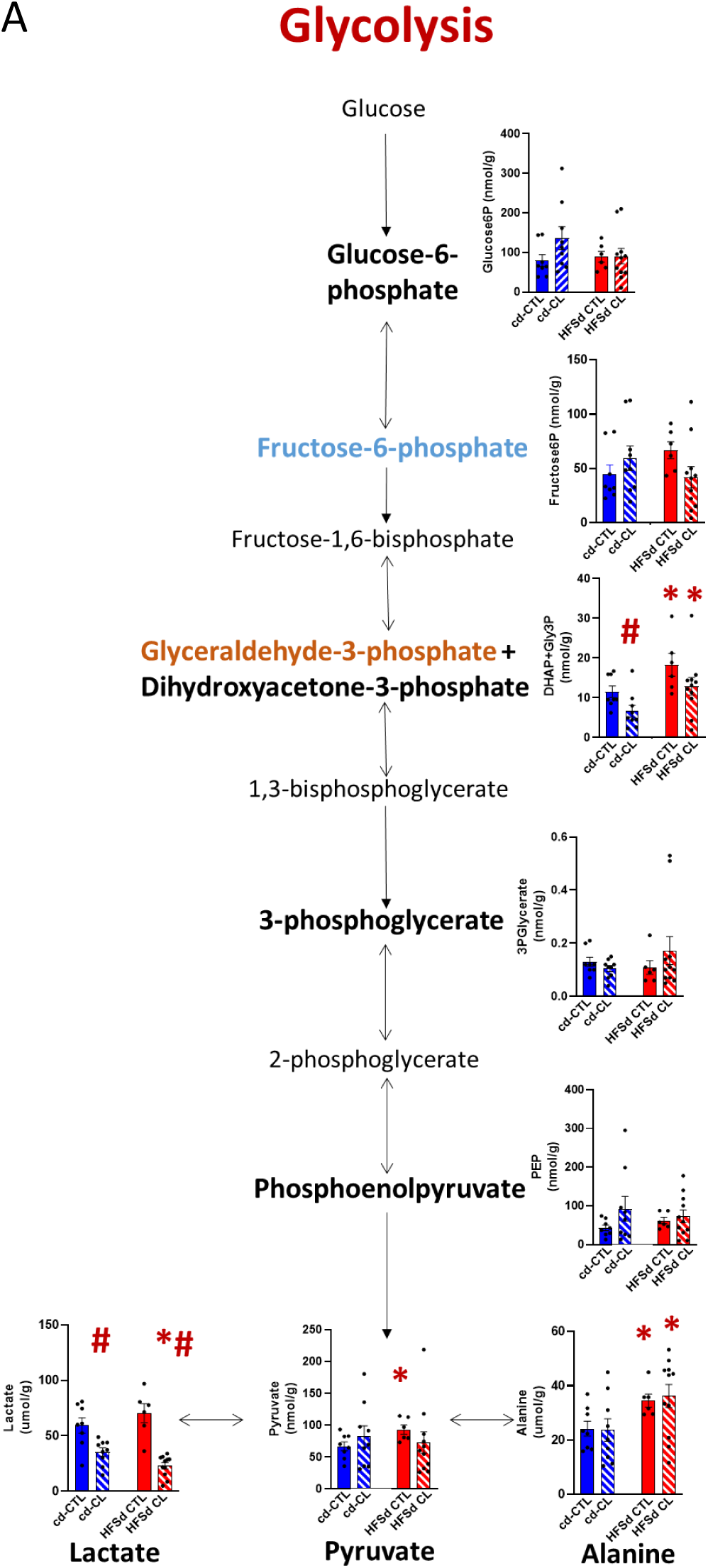
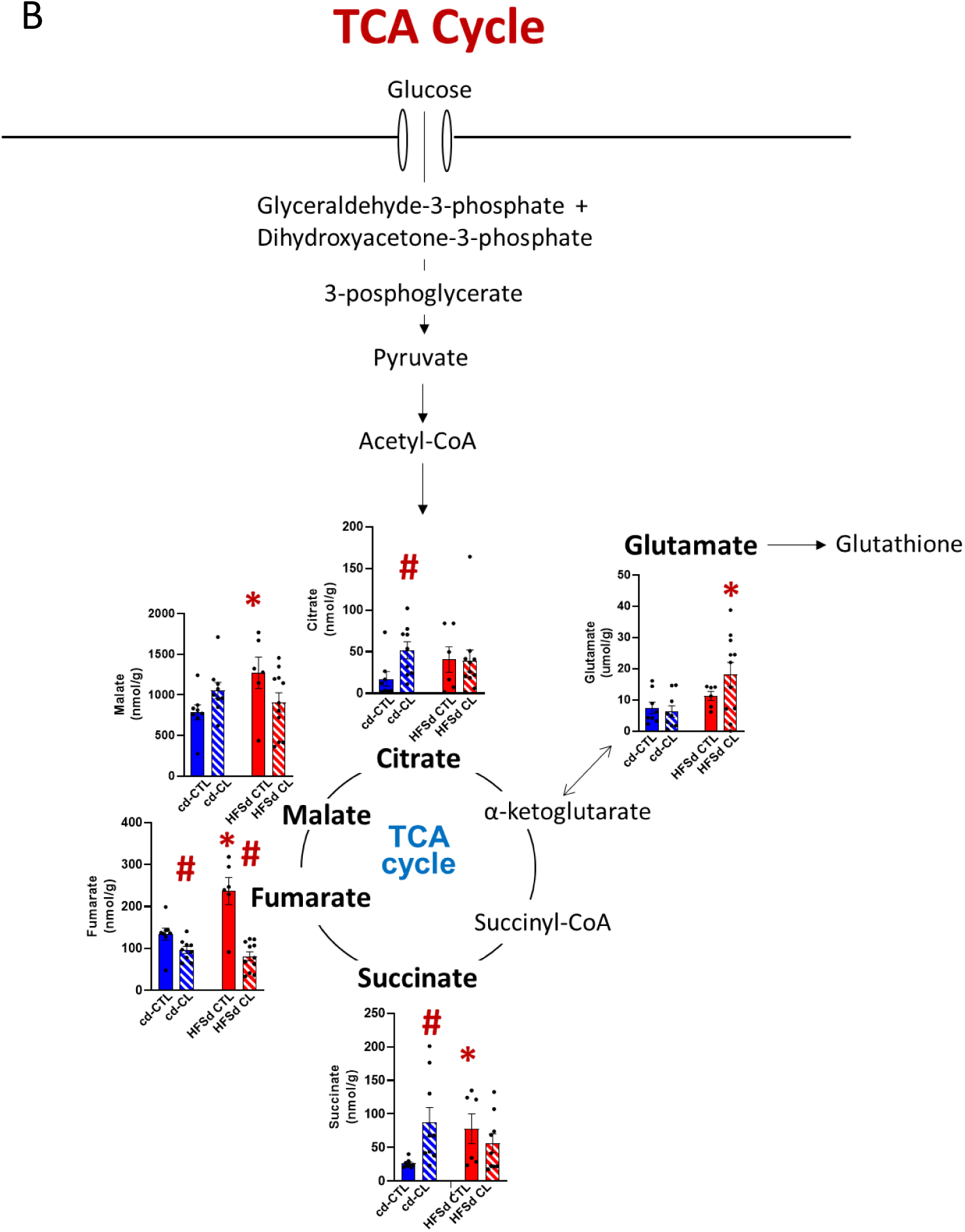
Metabolic analysis in heart in chow- and high-fat-sucrose-fed mice treated or not with CL316,243. Schematic representation of metabolic pathways from (A) glycolysis and; (B) oxidative phosphorylation with bar graph quantification of specific metabolites. *p-value ≤ 0.05 CD (chow diet, blue) *vs* HF-S (high-fat-sucrose, red), # p-value ≤ 0.05 CTL (control) *vs* CL (hatched).

Serine-to-Glycine ratio is increasingly recognized as predictor of cardiovascular risk, and Glycine is involved in key reactions of 1-Carbon metabolism (43). Thus, we expanded the modelization in that direction, as illustrated in Figure 10.

The model predicted significant changes in 1C-metabolism under CL treatment that resulted in increased SAM to SAH conversion, with increased donation of methyl groups (e.g., for DNA methylation); conversely, purine and pyrimidine metabolism (DNA synthesis) appear downregulated. Note that GSMM predicts increased conversion of Serine to Glycine, which is confirmed from our metabolomic measurements (suppl. Figure 5), and this could favour antioxidant glutathione synthesis from Glycine from several pathways, e.g., Glycine is a well-known precursor for glutathione synthesis through glutathione synthase.

Given the importance of lipid metabolism under HF-S, we expanded the analysis starting from the DEG coding enzymes involved in lipid metabolism in hearts from HF-S+/-CL. The most significant differentially expressed genes are illustrated in suppl. Table 3. Modelization of metabolism involving these genes predicted potentially important alterations by CL of sterol metabolism in the direction of reduced cholesterol synthesis (by Hmgcr), increased export (ABCG1) or storage (DGAT2, PLIN, ApoE), as well as production of oxysterols, which are known ligands of the transcription factor LXR (suppl. Figure 6).

To further support GSMMs results, we conducted additional targeted metabolomic measurements from cardiac tissue from the same mice in order to strengthen these observations (Suppl Fig 5). HF-S diet feeding was associated with increases in some intermediates of the glycolysis and oxidative metabolism, e.g.; increased levels of pyruvate and tricarboxylic acid (TCA) cycle intermediates (succinate, fumarate, malate). These increases were maintained under CL treatment (except a selective decrease in fumarate). The most notable change under CL was a strong decrease in lactate, both under chow and HF-S diets (Figure 11A), consistent with enhanced mitochondrial oxidative metabolism in the heart under CL. In addition, as indicated above, metabolomic analysis confirmed an increase in Glycine, strengthening its availability for 1-C metabolism whereas its preservation may indicate less need for antioxidant glutathione synthesis under CL.

To identify potential endocrine signals mediating communication between AT and heart, we used the epididymal AT transcriptomic data and filtered the CL-differentially expressed genes to retain those coding for secreted proteins. This analysis identified 198 candidate genes, including several molecules previously described as adipokines or circulating signalling factors, such as VEGFA, VEGFB, NRG4, NAMPT, or BDNF. We next used the nichenetr R package (v2.2.1.1) (44) to score those most likely to account for the transcriptomic changes measured in cardiac tissue, including genes accounting for the metabolic perturbations described above. The results, illustrated in Figure 12 identify the top-scored candidate mediators in AT, either up- (left panel) or down-regulated (right panel), and the inferred most likely associated target transcripts in the heart. Notably, all the up-regulated AT messengers act on target transcripts in the heart likely to be associated with preserved oxidative metabolism (e.g.; *Gadd45a*, *Igfbp3*, *Selenop*); conversely, all down-regulated AT messengers (e.g.; *Tgfb3*, *Ltgfb1*) are associated with cardiac transcripts involved in tissue remodeling (*Egr1*), cell cycle/proliferation (*Cdca3*, *Cd2*), including fibrosis.

**Figure 12:**
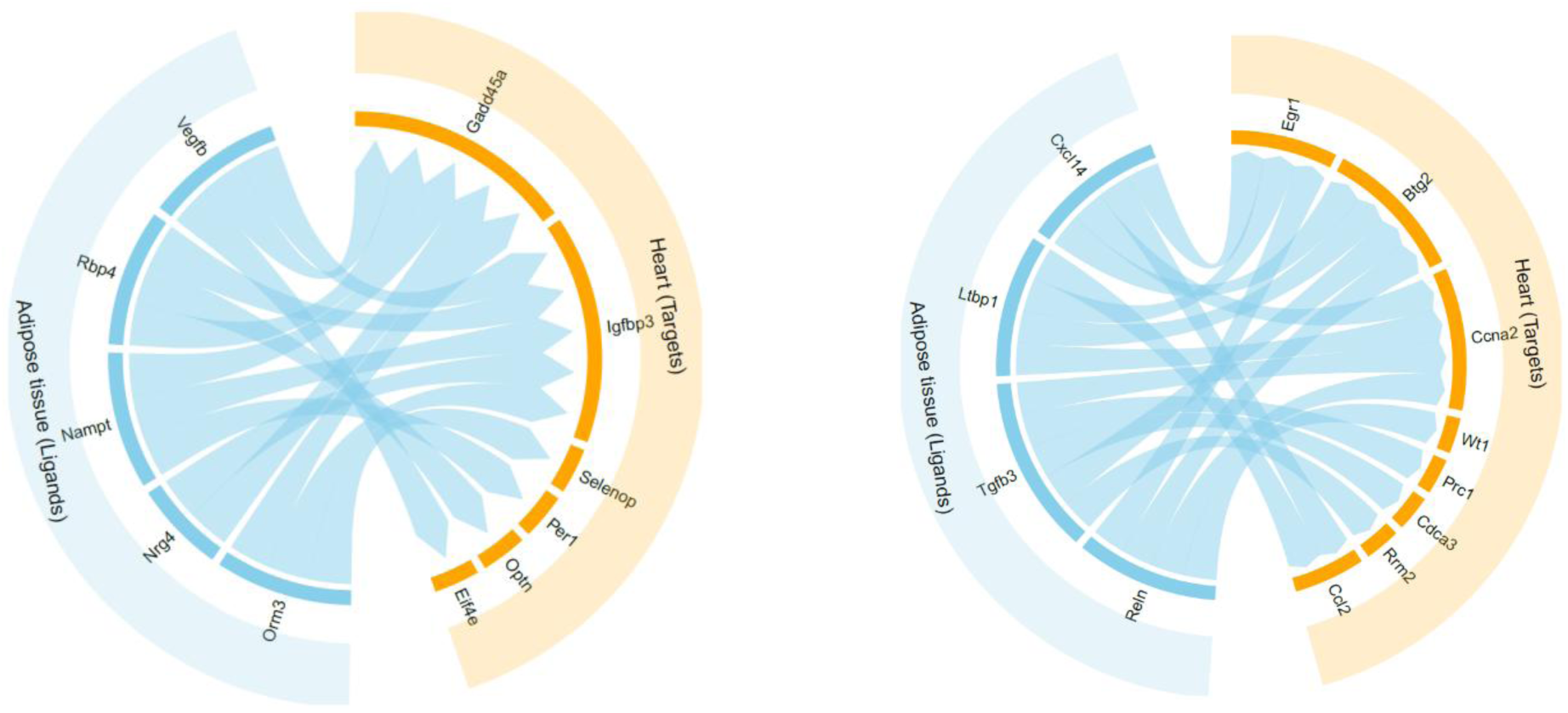
Prediction of putative endocrine/paracrine messengers mediating the adipocyte-cardiac cross-talk. Top upregulated (Left) or downregulated (Right) transcripts in adipose tissue (Blue) and corresponding upregulated (Left) or downregulated (Right) genes in cardiac tissue (Yellow), based on scores calculated using nichenetr R package (v2.2.1.1) from the transcriptome of both tissues.

## Discussion

In this study, we demonstrate that adipocyte β3AR plays a major role in mediating the systemic and cardiac effects of β3AR stimulation in the context of HF-S-induced obesity. Pharmacological activation of β3AR by CL316,243 (CL), a highly selective β3AR agonist in rodents, reduced body weight gain, improved glucose and insulin homeostasis, and preserved cardiac structure in obese mice. These beneficial effects were completely abolished in mice lacking β3AR specifically in adipocytes (B3 adipoKO), highlighting β3AR expressed in AT as a critical mediator of systemic metabolic health and cardioprotection. Mechanistically, the absence of β3AR in adipocytes prevented the CL-induced thermogenic activation of visceral AT (EPI AT) in obese mice, suggesting that defective adipose remodeling contributes to the loss of cardiac benefits. Bioinformatic analysis of combined transcriptomics and metabolomics in AT and cardiac extracts identified CL-altered metabolic pathways and candidate mediators most likely involved in this β3AR-dependent cross-talk.

BAT activation by cold exposure or by pharmacological stimulation has been extensively associated with improved metabolic health, through increasing energy expenditure and enhancing glucose and lipid utilisation, in both clinical and preclinical studies (24,45). In addition to its metabolic role, accumulating evidence indicates that it exerts remote cardiometabolic effects through UCP1-dependent thermogenesis and the secretion of bioactive adipokines or lipid mediators. Experimental studies have shown that BAT activation can protect against atherosclerosis (46) and attenuate pathological cardiac hypertrophy and fibrosis (47). A recent study showed that the adipocyte-specific genetic deletion of PRDM16 impairs the protective effects of perivascular AT against hypertension and arterial stiffness, highlighting beige adipocytes as modulators of vascular function (48). However, the cardiometabolic impact of β3AR-induced beige adipocyte activation during obesity remains poorly explored. Recently, Jousma *et al.* reported that enhancing (genetically) thermogenic AT activity improves cardiac function and lipid metabolism in a preclinical model of HFpEF (32). In the present study, we specifically investigated the cardiac impact of adipocyte β3AR stimulation, combined with unbiased transcriptomic and metabolomic approaches to explore the mechanisms underlying AT-heart communication.

We relied on a preclinical model that had been carefully established and validated in our previous work (41). In that study, we systematically evaluated two mouse strains and two obesogenic diets to identify the combination that most faithfully reproduces the key features of obesity-induced cardiac remodeling. This approach allowed us to select a robust and physiologically relevant model as the experimental basis for the present investigation. Using this model, we both induced “beiging” of AT with a β3AR agonist and generated an adipocyte-specific deletion of β3AR. The use of an inducible strategy was critical to avoid the compensatory adaptations that may occur in constitutive knockout models. The efficiency of β3AR deletion was verified at the genomic, transcriptomic and protein levels. Importantly, we verified that, in C57Bl6/J WT mice, β3AR is virtually undetectable in cardiac tissue, as previously reported, ruling out direct effects of the agonist in the heart. Finally, telemetric monitoring indicated that CL administration did not induce detectable β1AR-related cardiovascular side effects in our experimental conditions (i.e.; no significant increase in heart rate or mean blood pressure), supporting the specificity and safety of the pharmacological stimulation used in this study.

### Adipocyte β3AR is required for the systemic metabolic effects of CL

Consistent with previous reports (49,50), chronic activation of β3AR by CL promoted weight loss and improved insulin sensitivity in WT mice fed a HF-S diet. These effects were associated with a marked induction of thermogenic and beige markers specifically in epididymal AT, reflecting an improved metabolic activity. However, these metabolic benefits were completely lost in B3 adipoKO mice, which displayed weight gain, glucose intolerance and insulin resistance comparable to untreated HF-S controls. These findings demonstrate that β3AR expressed in adipocytes is essential for the metabolic response to β3AR agonist and that its absence is not compensated by β3AR signalling in other tissues.

### Loss of adipocyte β3AR abrogates β3AR-mediated cardioprotection

The major finding of this study is that the cardioprotective effect of β3AR activation is dependent on the presence of adipocyte β3AR. In WT obese mice, CL treatment protected the heart from the development of cardiac hypertrophy and LV interstitial fibrosis, confirming the beneficial impact of systemic β3AR stimulation on the heart. In contrast, B3 adipoKO mice exhibited concentric hypertrophy and increased cardiac mass similar to untreated HF-S controls, despite systemic CL administration. These results demonstrate that adipocyte β3AR is not only essential for systemic metabolic regulation but also exert remote protective effects on the heart.

This result should be distinguished, but also integrated with our previous studies that established the cardiac β3AR as an intrinsic cardioprotective modulator that counterbalances maladaptive β-adrenergic signalling. Specifically, we showed that direct cardiac β3AR activation promotes NO-dependent NRF2 activation, which enhances antioxidant defences through increased pentose-phosphate pathway metabolism during haemodynamic stress (51). Notably, in this previous study, we had used a model with cardiac myocyte-specific transgenic expression of the human β3AR, to reproduce its upregulation in human heart failure (52). In contrast, in the present non-failing model, endogenous cardiac β3AR expression was almost undetectable, as previously observed in mouse cardiac myocytes (42), and much lower compared with that observed in adipocytes, as confirmed by qPCR analysis. Nevertheless, β3AR is also expressed on the human and mouse endothelium where it promotes vasodilation (53) and thus may influence myocardial homeostasis through paracrine mechanisms. However, the loss of cardiac protection in our B3 adipoKO mice indicates that, in HF-S-induced obesity, β3AR-induced adipose-heart crosstalk plays a central role in maintaining cardiac health under metabolic stress.

### Temporal dissociation between metabolic and cardiac effects

An intriguing observation in our study is the temporal dissociation between the metabolic and cardiac effects of CL. While CL preserved insulin sensitivity during the first month of CL infusion, these metabolic benefits largely disappeared directly after treatment cessation. In contrast, the cardiac protection persisted after CL interruption because even after four months, we still observed protection against heart remodeling. This divergence suggests that the mechanisms driving metabolic improvements (e.g.; lipolysis, insulin sensitisation) may be transitory while the pathways underlying cardioprotection could involve more durable adaptations, possibly through (epi)genetic changes in cardiac cells, as suggested from our transcriptomic analysis (see below).

However, the heart-protecting effects of CL progressively declined over time during chronic treatment. This attenuation has been observed in several prolonged β3AR stimulation studies and may reflect adaptive desensitisation of β3AR signalling pathways during chronic obesity, a process known as catecholamine resistance (54,55). Several mechanisms have been proposed, including EPAC-mediated TRIB1 inhibition of *Adrb3* transcription (54), and IL1-beta-mediated activation of *Pde3a* to downregulate cAMP signalling. The latter identified neutrophils as the source of IL1beta to reduce β3AR-mediated lipolysis (56); this was proposed to preserve energy stores in the face of stress. These observations underscore the complexity of β3AR-mediated metabolic regulation and the potential need for combined (i.e., counteracting resistance mechanisms) or intermittent therapeutic strategies to sustain metabolic and cardiac benefits.

### Adipose-heart crosstalk as a mechanism of cardioprotection

The present findings reinforce the concept that AT is an active endocrine organ capable of modulating cardiac metabolism through the release of bioactive factors such as adipokines, cytokines, lipids and miRNA (13). The observation that CL-treated WT obese mice displayed preserved thermogenic gene profile in epididymal AT, whereas B3 adipoKO mice did not, supports the hypothesis that adipocyte β3AR signalling contributes to a favourable adipokine profile that may protect the heart from obesity-induced remodeling.

The localised specificity of the thermogenic response is also remarkable. CL induced the expression of *Ucp1*, *Cidea*, *Cox8b*, and *Elovl3* selectively in epididymal fat of obese mice, while the effect was not present in subcutaneous depots. This depot-specific pattern suggests that β3AR-dependent communication with the heart may differ between fat subtypes and the lack of beiging in EPI of B3 adipoKO mice may account, in part, for their loss of cardiac protection under HF-S. Notably, the epicardial fat shares a common embryologic origin with visceral AT (including EPI), so that similar CL-induced effects in epicardial AT may directly affect the underlying myocardium (57). However, it is important to note that human adipose depots do not map perfectly those of mice. Epididymal fat pad is often used as a surrogate for visceral fat, but humans lack a direct anatomical equivalent of this depot (58). Therefore, while our mouse data provide strong mechanistic insight, caution is warranted when investigating the depot-specific β3AR effects observed here to human physiology.

The integrative transcriptomic and metabolomic analyses provide mechanistic insights into the β3AR-mediated adipose-heart communication underlying β3AR-mediated cardioprotection.

In the AT, we showed that CL induces metabolic and phenotypic changes in the EPI AT of obese mice toward an increased oxidative metabolism, reflecting active thermogenic process, accompanied by reduced inflammatory, hypoxic and EMT signatures. These changes are indicative of improved AT function and remodeling. In line with this context, several secreted factors identified in our adipose DEG dataset are known to regulate local AT homeostasis.

Among these, VEGFA (vascular endothelial growth factor A), and to a lesser extent VEGFB, are key regulators of angiogenesis, a critical process for maintaining adequate oxygen and nutrient supply during AT expansion. Adequate vascularization is essential to prevent hypoxia, inflammation and metabolic dysfunction. In addition to their vascular effects, VEGF signalling has been shown to promote metabolic health in AT by protecting against diet-induced obesity and insulin resistance (59). Moreover, VEGFA expression has been associated with the induction of AT browning and thermogenesis (60), which are consistent with our observations. Notably, despite increased expression of VEGF ligands, our hallmark analysis revealed a decrease in angiogenesis-related pathways in EPI AT following CL treatment. In obesity, angiogenesis typically increases as a compensatory response to hypoxia, although this response is often insufficient to fully restore tissue oxygenation. The reduction in angiogenic signature observed here may likely reflect an improvement in AT function. Indeed, the concomitant downregulation of hypoxia and inflammation-related pathways suggests a reduced need for active vascular remodeling, consistent with a shift toward a healthier and better-oxygenated AT.

On the cardiac side, our transcriptomic data are consistent with previous studies of HF-S-induced cardiac remodeling, which report DEG predominantly enriched in pathways related to fatty acid/PPAR signalling, glucose metabolism and mitochondrial processes (61). By overlaying these with transcripts differentially expressed in HF-S hearts under CL, we identified 43 genes reciprocally regulated between HF-S and HF-S+CL (see Venn diagram in Figure 7F and supplemental Table 2), most of which are involved in cardiac metabolism. Using these as “seed” genes, we reconstructed metabolic pathways using the mouse generic Genome Scale Metabolic Modeling (62). Based on several assumptions, this approach uses gene expression data to infer metabolic fluxes with a pathway reaction score (PRS) that predicts its up- or downregulation. The results predict a reinforcement of oxidative metabolism through the Krebs cycle in HF-S+CL hearts, through increased metabolism of glutamate and glycolysis-derived pyruvate, at the expense of lactate accumulation. Importantly, the latter was verified experimentally through targeted metabolomic analysis from the same heart extracts. This observation would be in line with an adaptative selection of glucose as substrate for oxidation in the face of excessive fat under obesogenic diet (63). The model also predicts a decrease in Serine synthesis from glycolytic intermediates; while Serine levels are unchanged under CL in our metabolomic measurements, Glycine is increased, which is usually associated with an improvement in metabolic profile (64). Further modelization of Glycine metabolism predicts increased fluxes in 1-carbon metabolism towards methylation reactions rather than purine/pyrimidine synthesis. This would suggest restrained DNA synthesis for cell division/proliferation, compatible with decreased fibrosis while DNA methylation would adaptively maintain stability in gene expression regulation. The preservation of Glycine available for glutathione synthesis may be indicative of reduced demand compatible with reduced oxidant stress under CL.

As high-fat feeding profoundly influences cardiac lipid metabolism, we further expanded our DEG-based metabolic modelization from cardiac transcripts most significantly affected under CL treatment. Most genes upregulated under CL are associated with enzymatic reactions involved in omega-3, −6 (*Hacd4*), leukotriene and prostaglandin synthesis (*Hpgds*) and cholesterol disposition (*ApoE*, *Plin*, *Cyp27a1*, *Dgat2*, *Cyp4v3*). Omega-3, −6 synthesis may be important for the preservation of cardiolipin in the face of increased mitochondrial turnover, and some prostaglandins are known to be involved as protective lipid mediators. Notably, modelization of cholesterol metabolism indicates downregulation of cholesterol synthesis (*Hmgcr*) or upregulation of enzymatic reactions associated with either cholesterol export (*Abca1*), storage in lipid droplets (*Plin3*, *Dgat2*, *Apoe*) or oxidation to oxysterols (*Cyp27a1*). As free intracellular cholesterol is toxic, these reactions would favour protection both from endogenous cholesterol synthesis and import from lipoproteins. Notably, oxysterol are ligands of LXR that, together with other transcriptional co-regulators (e.g., RXR) are known to drive the expression of a number of genes with protective effects against oxidative stress, thereby preserving mitochondrial integrity and metabolism (65–67).

One strength of our combined analysis of AT and cardiac transcriptome is the identification of the most likely candidate mediators for the inter-organ cross-talk using the nichenetr R package (v2.2.1.1) (68). Overlaying our CL-induced DEG with curated databases allows to associate protein-coding genes in the donor (AT) tissue with genes and signalling pathways most represented in the recipient (heart) tissue. Based on a probability score, the results identify both upregulated and downregulated messengers associated with pathways remarkably relevant to our observed phenotypes, including metabolic predictions.

Among these, neuregulin 4 (NRG4) emerges as a particularly interesting mediator. NRG4 is an adipokine predominantly produced by thermogenic adipocytes and has been reported to be implicated in systemic metabolic health and insulin sensitivity (69). In addition, emerging evidence suggests that NRG4 can exert cardioprotective effects by activating ErbB signalling pathways, limiting pathological cardiac remodeling including hypertrophy and fibrosis, in metabolic diseases (70,71). In addition, NRG4 has been associated with liver protection during MAFLD (metabolic dysfunction-associated steatotic liver disease) (72,73). Its upregulation under CL is compatible with the predicted beneficial effects on myocardial metabolism (improved glucose uptake and oxidation), as well as observed attenuated cardiac remodeling.

NAMPT (nicotinamide phosphoribosyltransferase) a key enzyme in NAD⁺ biosynthesis, plays a central role in cellular energy homeostasis. NAD+ (nicotinamide adenine dinucleotide) is an essential co-factor involved in multiple metabolic processes such as glycolysis, FAO, oxidative phosphorylation and TCA cycle. In addition, it regulates mitochondrial function through protein deacetylation via sirtuins (74,75). Importantly, beyond its intracellular role, NAMPT exists in a secreted form (extracellular NAMPT, also referred as visfatin), which can be released by the AT and transported systemically within extracellular vesicles. Circulating NAMPT has been shown to promote NAD+ biosynthesis in peripheral tissues and to contribute to pancreas beta cell function, insulin secretion and protects against age-related decline in mice via SIRT-dependent pathways (76–78). Although circulating NAMPT has been correlated with improved cardiac function (79,80), a direct link between AT-derived NAMPT and heart has not yet been formally demonstrated. Signalling by NAMPT is again compatible with our predicted improved cardiac metabolic capacity, potentially linked to enhanced systemic NAD⁺ availability.

Members of the VEGF family may also contribute to the observed cardiac transcriptional changes. Vegf-b is known to increase cardiac vascularization and promote glucose uptake and oxidation (81,82), and its signalling has been associated with *Pcx*, *Abat* and *Kyat3*, in line with our metabolic predictions. Cardiac adenoviral transduction of Vegfb also reverses adverse myocardial remodeling in aging mice (83). Orm3 has been implicated in antioxidant defense through Nrf2 (84), while the vitA binding protein, Rbp4 plays a dual role as adipokine that is context-dependent, even worsening cardiac remodeling in the context of diabetes (85,86). Notably, most of these candidate mediators are associated with upregulation of *Gadd45a* (growth arrest and DNA damage inducible 45a) in the cardiac transcriptome, which has been implicated in cardiac protection from inflammation and fibrosis (87). Likewise, *Igfbp3* has been implicated in the preservation of mitochondrial integrity and respiration (88).

Among other cardiac target genes, *Selenop* (selenoprotein P) also contributes to the regulation of cardiac redox homeostasis. As a selenium transport protein, Selenop supports the activity of antioxidant enzymes such as glutathione peroxidases, which limit oxidative stress (89). *Optn* (Optineurin) promotes the elimination of damaged mitochondria through mitophagy (90), while the regulator of protein synthesis, *Eif4e* increases the translation of RNA coding for mitogenesis (91). Of interest, among the highly-ranked cardiac transcripts, *Per1* (Periodic circadian regulator 1), a negative transcriptional regulator of the CLOCK gene, mediates the circadian control of metabolism, a key controller of metabolic flexibility that is lost under high fat diet (92).

On the downregulation side, *Tgfb3* (Tgf-beta3) and *Ltgfb1* (Latent TGF-beta binding protein-1) indicate reduced pro-fibrotic signalling under CL, as observed in our cardiac phenotype, while decreased *Cxcl14* would result in decreased recruitment of inflammatory cells and *Reln* (Reelin) in decreased endothelial proliferation. Associated cardiac transcripts include downregulation of the chemokine *Ccl2*, compatible with reduced inflammation, while *Cdca3* (cell division cycle-associated protein 3, a binding protein associated with cell division), *Ccna2* (Cyclin A2, activator of cyclin-dependent kinase 2) both indicate reduction of cell division/proliferation, together with downregulation of other regulators of cell cycle (*Btg2*), cytokinesis (*Prc1*), DNA synthesis (*Rrm2*) or RNA processing (*Wt1*). Notably, some of the downregulated mediators are associated with downregulated *Egr1*, a master regulator of the cardiac « fetal gene program », compatible with preservation from adverse cardiac remodeling by CL (93,94).

Altogether, these findings support a model in which β3-adrenergic activation of AT promotes the secretion of multiple signalling factors capable of acting locally within adipose depots, and systemically, by exerting endocrine effects on the heart. These adipose-derived signals may contribute to the restoration of metabolic pathways, modulation of inflammatory responses and overall improvement of myocardial homeostasis in the context of diet-induced obesity.

### Limitations

Our study was conducted exclusively in male mice in order to eliminate sex-related variables. Therefore, we cannot exclude sex-specific differences in β3AR signalling and adipose-cardiac interactions. While we carefully examined several metabolically distinct AT types, other adipose depots may contribute to the observed effects and warrant investigation. Epicardial AT, in particular, has gained attention for its role in cardiac health due to its proximity to the myocardium. However, its limited quantity in mice did not allow a comprehensive analysis in our model. While our metabolomic measurements support, in part, the fluxes predicted from GSMM, our bioinformatic approach has inherent limitations. The results, however, are fully in line with known pathways activated by our candidate mediators. While we did not perform measurements of cardiac lipidomics, these mainly reflect accumulation of lipotoxic intermediates and changes in membrane composition, while metabolomic measurements are closer to changes in energy metabolism which is the focus of our study.

## Conclusion

Altogether, our findings establish for the first time a functional link between adipocyte β3AR activation and a protection from obesity-induced cardiac remodeling. We demonstrate that adipocyte β3AR is a critical mediator of the systemic metabolic and cardiac benefits induced by chronic β3AR stimulation in mice. By promoting thermogenic activation of epididymal fat, adipocyte β3AR improved metabolic homeostasis and prevented the development of cardiac hypertrophy under obesogenic conditions. Conversely, its absence abolishes CL-induced metabolic improvements, epididymal AT beiging and cardioprotection, underscoring the importance of adipose-heart communication during metabolic diseases. This study raises the possibility to use β3AR signalling as a potential therapeutic target to mitigate obesity-associated CVDs. However, the transient efficacy observed with prolonged stimulation suggests that it will likely require combinatorial or adaptive strategies to reverse catecholamine resistance (54,55).

## Grant acknowledgments

This work was funded by a grant (PDR T.0008.22) from the Fonds National de la Recherche Scientifique (FNRS). LC is a recipent of a Fellowship (Aspirante) of the FNRS.

AG received grant support from European Union’s Horizon Europe Research and Innovation Programme for the project “PAS GRAS: De-risking metabolic, environmental and behavioral determinants of obesity in children, adolescents and young adults” under grant agreement No. 101080329.

and

European Union’s Horizon Europe research and innovation programme for the project “GRIPonMASH: Global Research Initiative for Patient screening on MASH.” The GRIPonMASH project is supported by the Innovative Health Initiative Joint Undertaking (IHI JU) under grant agreement No 101132946.

## Author contributions

LC, LM, HE, DDM, AM, CB, SP, FC: data acquisition, data analysis, manuscript editing; SP, FC, SS, AG: metabolomics and modelisation, manuscript editing; AL, JA, LG: transcriptomics and bioinformatic analysis, manuscript editing; FC: metabolomics profiling and interpretation of the results; SP: metabolomics profiling and data analysis; SS: genome-scale metabolic modeling and interpretation of the results; AG: supervision of the metabolomics and modeling analyses and interpretation of the results; LM, CD, AG, JLB: conceptualisation, data analysis, manuscript editing; AG, JLB: funding acquisition.

