## Supplemental material for "Beige/brown fat-mediated cardiac protection from high-fat diet is dependent on adipocyte beta3-adrenergic receptor"

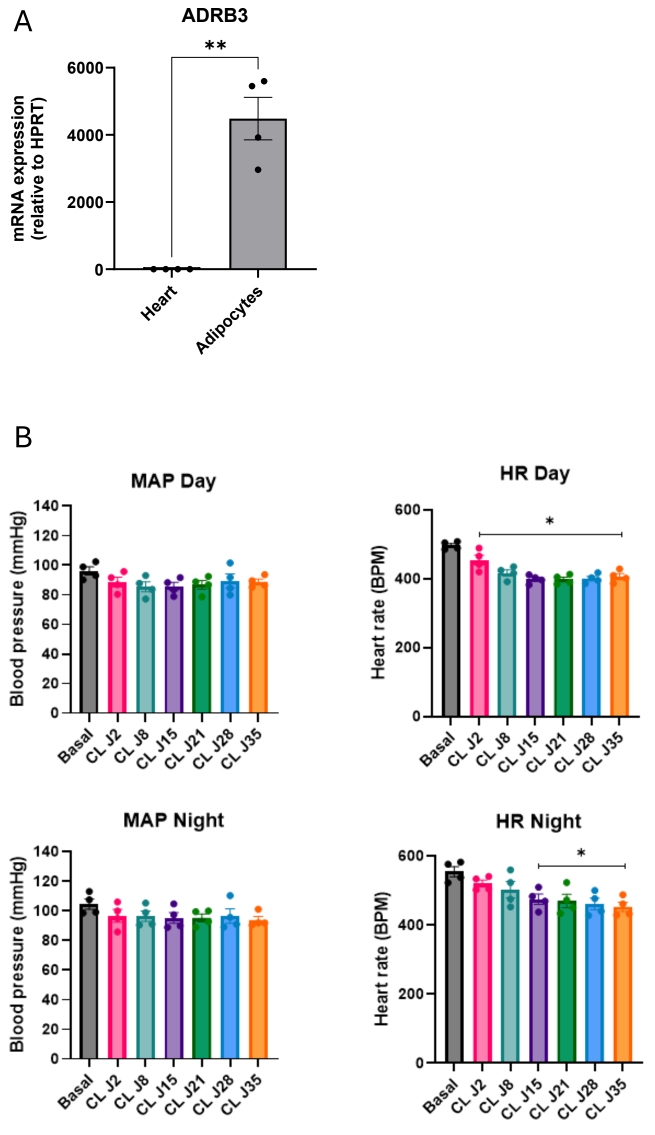


**Supplemental Figure 1:** (A) Relative mRNA expression of *Adrb3* from heart and isolated adipocytes from inguinal AT of chow diet mice; (B) Telemetric measurement of mean arterial pressure (MAP) and heart rate of mice during day (up) and night (down) at baseline and at 2, 8, 15, 21, 28 and 35 days of CL infusion through osmotic minipump. Data are presented as means +/- SEM (N = 4 per group) and analysed by Student's t-test or by 1-way ANOVA (*p<0,05, **p<0,01).

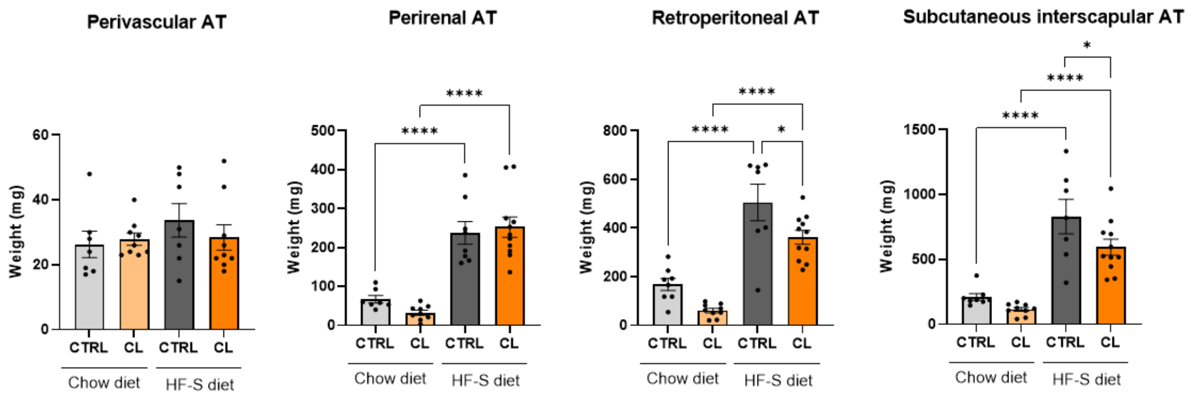

**Supplemental Figure 2:** Weight of perivascular, perirenal, retroperitoneal and subcutaneous interscapular AT after 24 weeks of diet, in chow- (light) and HF-S-fed (dark) mice treated (orange) or not (grey) with CL316,243 (CL). Data are presented as means +/- SEM (N = 6-9 per group) and analysed by 2-way ANOVA (*p<0,05, ****p<0,0001).

| | C57Bl6/J | | Chow diet | HF-S diet | Chow diet + CL | HF-S diet + CL |
|---|---|---|---|---|---|---|
| Weight (mg) | BAT | Interscapular | 219,4 ± 15,7 | 628,0 ± 42,6 | 120,4 ± 8,7 | 259,5 ± 22,1 |
| | Subcutaneous | Interscapular | 210,0 ± 25,1 | 830,1 ± 133,5 | 113,9 ± 14,8 | 594,5 ± 62,64 |
| | | Inguinal AT | 332,5 ± 33,1 | 1361,4 ± 137,1 | 222,1 ± 19,8 | 1471,3 ± 71,3 |
| | Visceral | Pericardial AT | 21,3 ± 3,9 | 66,3 ± 10,5 | 6,2 ± 1,4 | 44,2 ± 6,6 |
| | | Epididymal AT | 636,5 ± 62,7 | 1359,3 ± 78,3 | 240,9 ± 25,8 | 1363,0 ± 83,3 |
| | | Perirenal AT | 66,9 ± 9,4 | 237,4 ± 29,1 | 32,3 ± 6,2 | 252,5 ± 26,0 |
| | | Retroperitoneal AT | 168,0 ± 25,1 | 504,9 ± 75,2 | 60,7 ± 9,1 | 361,7 ± 28,3 |
| | | Mesenteric AT | 252,9 ± 40,5 | 643,0 ± 130,0 | 142,4 ± 31,1 | 766,4 ± 87,5 |
| | Other | Perivascular AT | 26,3 ± 4,1 | 33,7 ± 5,2 | 27,9 ± 1,9 | 28,4 ± 3,9 |

**Supplemental Table 1:** Weight (in milligrams) of the adipose tissues (AT) of the C57Bl/6J mice treated with chow diet (CD) or high-fat-sucrose (HF-S) diet with or without CL316,243 (CL) at 24 weeks of diets. Interscapular brown adipose tissue (BAT), inguinal, epididymal, perirenal, retroperitoneal AT were weighted on the right side of the mice. The mass of the pericardial, interscapular subcutaneous, perivascular and mesenteric AT are the total mass of the mice. Data indicate the mean +/- SEM (N = 7-9 mice per group).

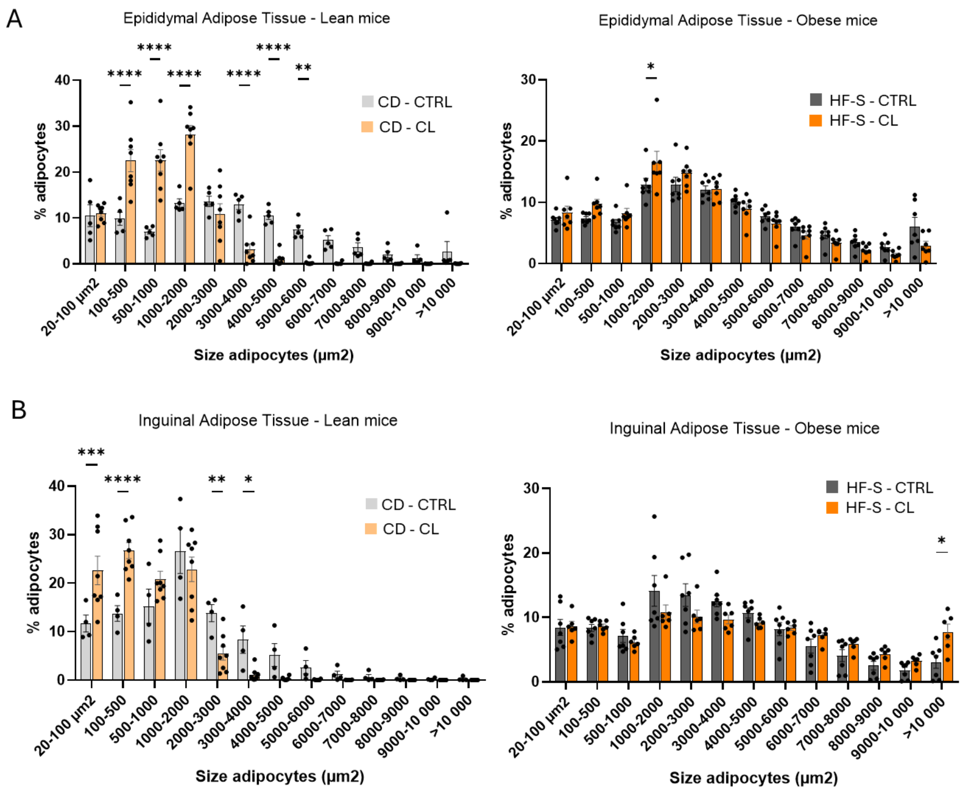


**Supplemental Figure 3:** Adipocyte size distribution (20-1000 µm²) in epididymal (A) and inguinal (B) AT of chow- (light) and high-fat-sucrose-fed (dark) mice treated (orange) or not (grey) with CL316,243 (CL), performed with haematoxylin-eosin (HE) histological labelling. Data are presented as means +/- SEM (N = 5-9 per group) and analysed by 2-way ANOVA (*p<0,05, **p<0,01, ***p< 0,001, ****p<0,0001).

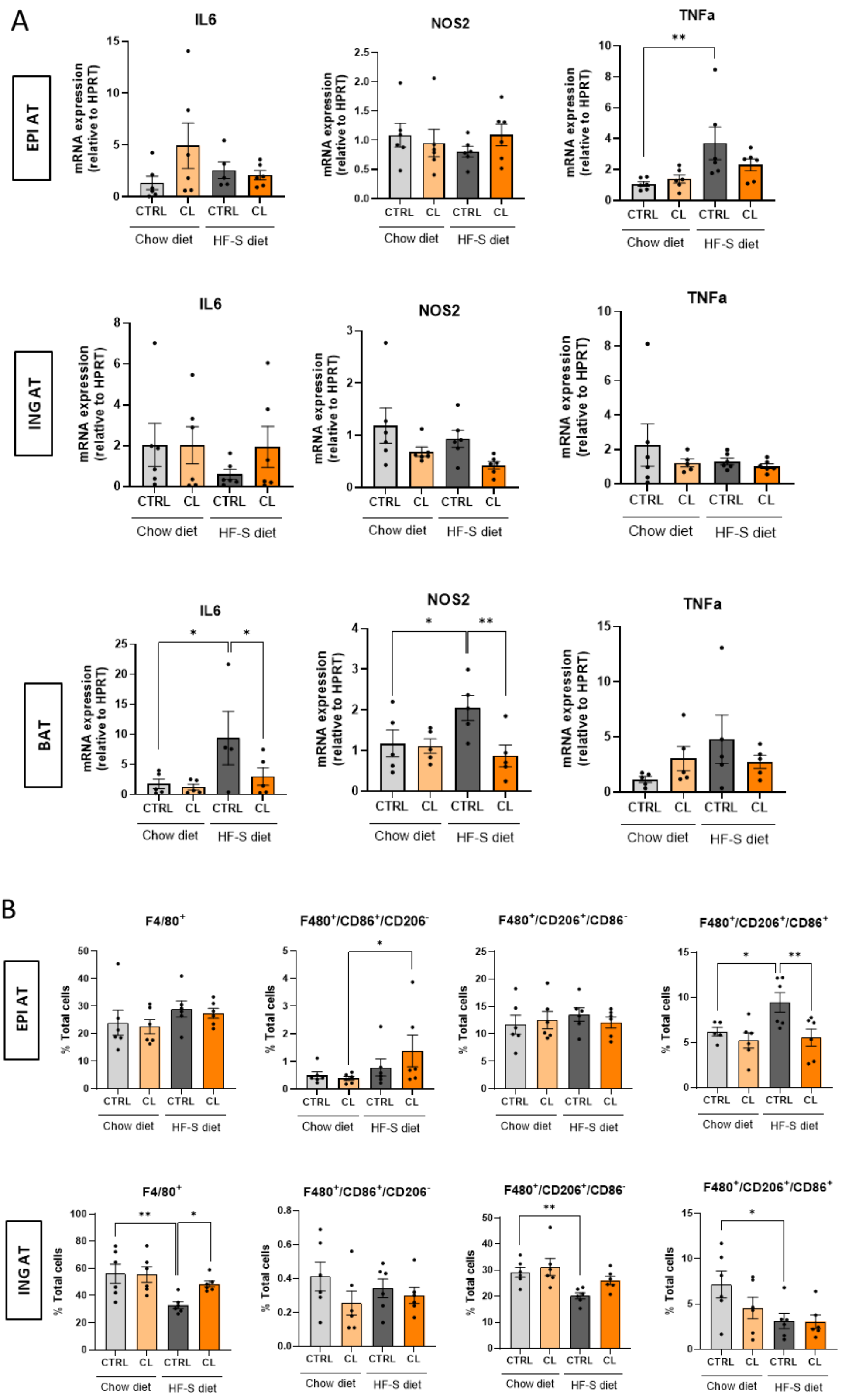


**Supplemental Figure 4:** (A) mRNA expression of inflammatory cytokines TNFα, NOS2 and IL6 in epididymal (EPI), inguinal (ING) and brown (BAT) adipose tissues (AT) of chow- (light) and HF-S-fed (dark) mice treated (orange) or not (grey) with CL316,243 (CL) after 24 weeks of diet; (B) proportion of total macrophages (F4/80+), CD86 positive macrophages (F4/80+/CD86+/CD206-), CD206 positive macrophages (F4/80+/CD206+/CD86-) and double CD86 and CD206 positive macrophages (F4/80+/CD206+/CD86+) by multiplex staining on EPI and ING AT after 24 weeks of diet. Data are presented as means +/- SEM (N = 6-9 per group) and analysed by 2-way ANOVA (*p<0,05, **p<0,01).

| ENSEMBL | Gene | CD *vs* HF-S diet - Heart | HF-S diet *vs* HF-S+CL - Heart | CD *vs* CD+CL - Heart |
|---|---|---|---|---|
| ENSMUSG00000047085 | Lrrc4b | DOWN | UP | ns |
| ENSMUSG00000057880 | Abat | DOWN | UP | ns |
| ENSMUSG00000027524 | Edn3 | UP | DOWN | ns |
| ENSMUSG00000079105 | C7 | DOWN | UP | UP |
| ENSMUSG00000004347 | Pde1c | DOWN | UP | ns |
| ENSMUSG00000030889 | Vwa3a | UP | DOWN | ns |
| ENSMUSG00000003545 | Fosb | UP | DOWN | ns |
| ENSMUSG00000020178 | Adora2a | UP | DOWN | ns |
| ENSMUSG00000035105 | Egln3 | UP | DOWN | ns |
| ENSMUSG00000028766 | Alpl | UP | DOWN | ns |
| ENSMUSG00000025221 | Kcnip2 | DOWN | UP | ns |
| ENSMUSG00000036006 | Ripor2 | DOWN | UP | ns |
| ENSMUSG00000027559 | Car3 | DOWN | UP | ns |
| ENSMUSG00000040213 | Kyat3 | DOWN | UP | ns |
| ENSMUSG00000028278 | Rragd | DOWN | UP | ns |
| ENSMUSG00000060548 | Tnfrsf19 | DOWN | UP | ns |
| ENSMUSG00000020051 | Pah | DOWN | UP | ns |
| ENSMUSG00000042613 | Pbxip1 | DOWN | UP | ns |
| ENSMUSG00000027030 | Stk39 | DOWN | UP | ns |
| ENSMUSG00000024892 | Pcx | DOWN | UP | ns |
| ENSMUSG00000019851 | Perp | DOWN | UP | ns |
| ENSMUSG00000045573 | Penk | DOWN | UP | ns |
| ENSMUSG00000005667 | Mthfd2 | UP | DOWN | ns |
| ENSMUSG00000026672 | Optn | DOWN | UP | ns |
| ENSMUSG00000050106 | Tmc8 | UP | DOWN | ns |
| ENSMUSG00000015112 | Slc25a13 | DOWN | UP | ns |
| ENSMUSG00000002771 | Grin2d | DOWN | UP | ns |
| ENSMUSG00000038418 | Egr1 | UP | DOWN | ns |
| ENSMUSG00000104736 | NA | DOWN | UP | ns |
| ENSMUSG00000018381 | Abi3 | UP | DOWN | ns |
| ENSMUSG00000046470 | Sox18 | UP | DOWN | ns |
| ENSMUSG00000025889 | Snca | UP | DOWN | ns |
| ENSMUSG00000055435 | Maf | DOWN | UP | ns |
| ENSMUSG00000023505 | Cdca3 | UP | DOWN | ns |
| ENSMUSG00000042351 | Grap2 | UP | DOWN | ns |
| ENSMUSG00000038943 | Prc1 | UP | DOWN | ns |
| ENSMUSG00000115125 | NA | DOWN | UP | ns |
| ENSMUSG00000032006 | Pdgfd | UP | DOWN | ns |
| ENSMUSG00000038801 | Scgb1c1 | DOWN | UP | UP |
| ENSMUSG00000016520 | Lnx2 | UP | DOWN | ns |
| ENSMUSG00000021250 | Fos | UP | DOWN | ns |
| ENSMUSG00000031461 | Myom2 | DOWN | UP | UP |
| ENSMUSG00000085779 | NA | DOWN | UP | ns |

**Supplemental Table 2:** List of the 43 genes reciprocally regulated in the heart by high-fat sucrose (HF-S) diet and CL treatment in HF-S-fed mice. Genes were identified by intersecting differentially expressed genes from the comparison between chow diet (CD) and HF-S diet hearts, and between HF-S diet and CL-treated HF-S hearts. The direction of regulation (up or down) in each condition is indicated. ns: non-significant.

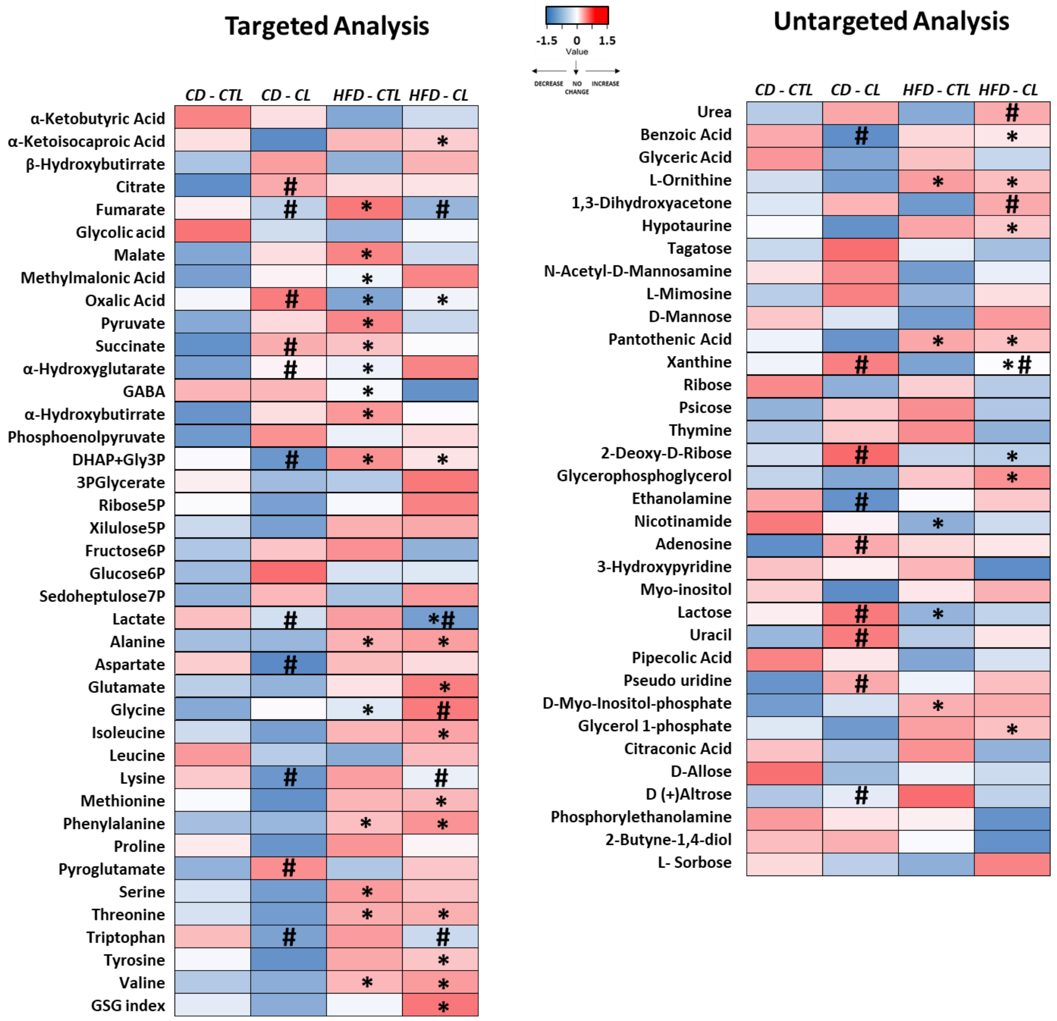


**Supplemental Figure 5:** Heatmap from targeted and untargeted metabolic analysis of cardiac tissue from chow- (CD) and HF-S (high-fat-sucrose)-fed mice treated or not with CL316,243 (N=8-9 per group). * p-value ≤ 0.05 CD *vs* HF-S, # p-value ≤ 0.05 CTL (control) *vs* CL.

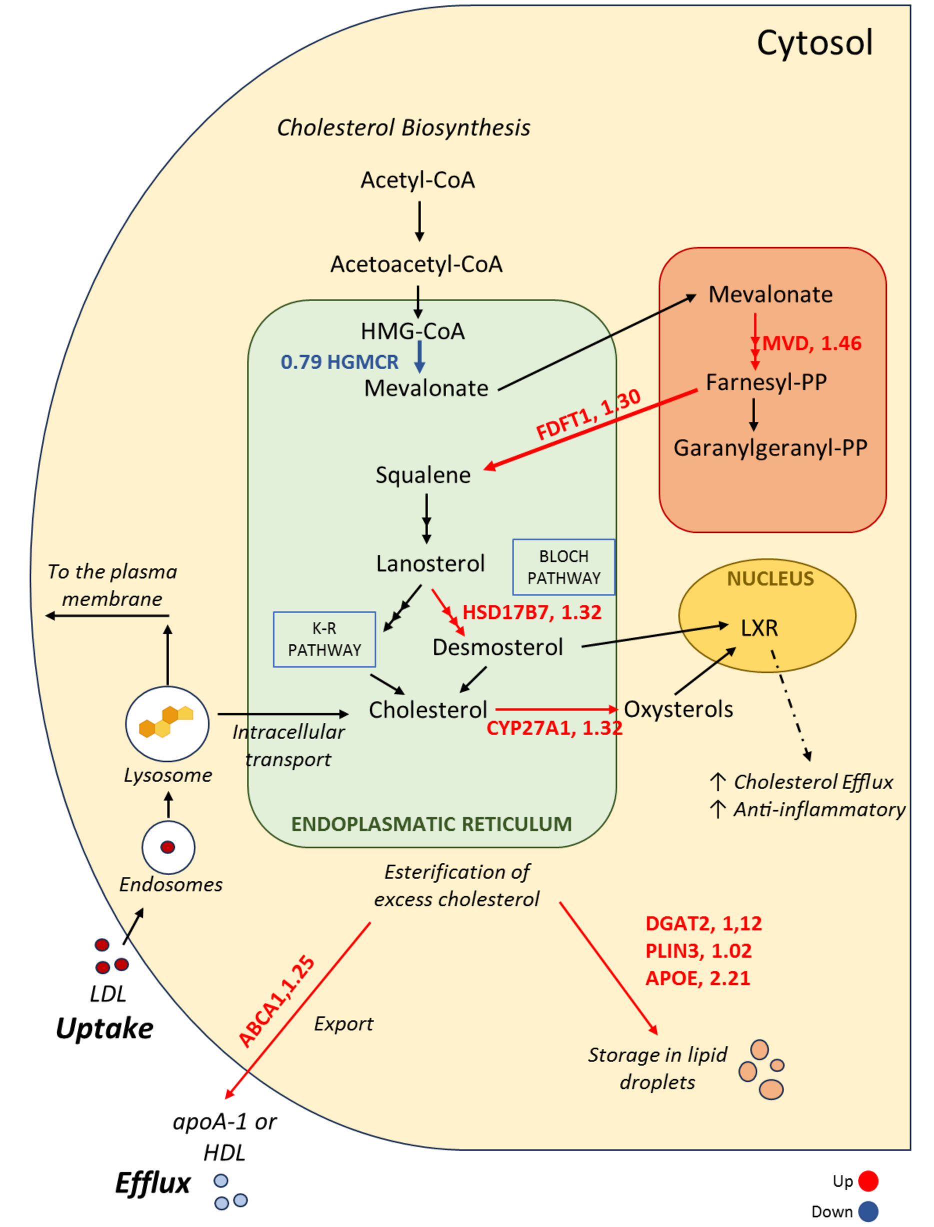


**Supplemental Figure 6:** Metabolic network illustrating pathways alterations observed in the cholesterol biosynthesis, metabolism and homeostasis subsystems in HF-S conditions with versus without CL. Reactions exhibiting changes in flux ratios greater than 1.25, lower than 0.9, or associated with differentially expressed genes are highlighted in red (upregulated) or blue (downregulated).

| status | external_gene_name | log2FoldChange | newfdr | AssociatedSubsystems |
|---|---|---|---|---|
| UP | Apoe | 1.14 | 1.29E-06 | Protein assembly |
| UP | Plin3 | 0.36 | 0.065268 | Pool reactions |
| UP | Cyp27a1 | 0.46 | 0.058956 | Bile acid biosynthesis; Cholesterol metabolism; Glycerophospholipid metabolism; Miscellaneous; Vitamin D metabolism |
| UP | Hacd4 | 0.83 | 0.016765 | Fatty acid biosynthesis (unsaturated); Fatty acid elongation (even-chain); Fatty acid elongation (odd-chain); Omega-3 fatty acid metabolism; Omega-6 fatty acid metabolism |
| DOWN | Alpl | -0.63 | 8.72E-05 | Biopterin metabolism; Glycerolipid metabolism; Glycerophospholipid metabolism; Miscellaneous; Xenobiotics metabolism |
| UP | Hpgds | 1.51 | 0.000185 | Arachidonic acid metabolism; Eicosanoid metabolism; Estrogen metabolism; Leukotriene metabolism; Phenylalanine, tyrosine and tryptophan biosynthesis; Prostaglandin biosynthesis |
| UP | Dgat2 | 0.35 | 0.076371 | Acylglycerides metabolism; Retinol metabolism |
| UP | Cyp4v3 | 0.83 | 0.065268 | Steroid metabolism; Vitamin D metabolism |

**Supplemental Table 3:** Genes related to lipid, fatty acids or sterols pathways significantly altered in HF-S +/- CL.
